# BOTany Methods: Accessible Automation for Plant Synthetic Biology

**DOI:** 10.1101/2025.08.21.671538

**Authors:** Moni Qiande, Abigail Lin, Lianna Larson, Cătălin Voiniciuc

## Abstract

Most members of the synthetic biology community, particularly plant scientists, lack access to liquid handling robots to scale up experiments, enhance reproducibility, and accelerate the Design, Build, Test, Learn cycle. Biofoundries enable high throughput data acquisition to train AI models and to develop new bioproducts, but they are capital-intensive to set up and not widely distributed. Entry-level, 3D-printed robots offer more affordable alternatives, but suffer from a shortage of validated protocols that can be modified without prior coding experience. To enhance access to biological automation, we developed a collection of modular BOTany Methods using Opentrons OT-2 robots to streamline the most common methods for molecular biology research and education. Our comprehensive workflow offers automation for a variety of procedures, ranging from simple but repetitive tasks (such as primer dilution and PCR setup) to more complex operations, including Plant Modular Cloning (MoClo), bacterial transformation, and plasmid extraction. Our BOTany Methods enable undergraduate students and other early career researchers to run designer experiments using table-based inputs, without editing the custom Python scripts. This pipeline enables end-to-end molecular cloning with minimal user intervention, enhancing throughput and traceability for synthetic biology applications.

**Graphical Abstract:** 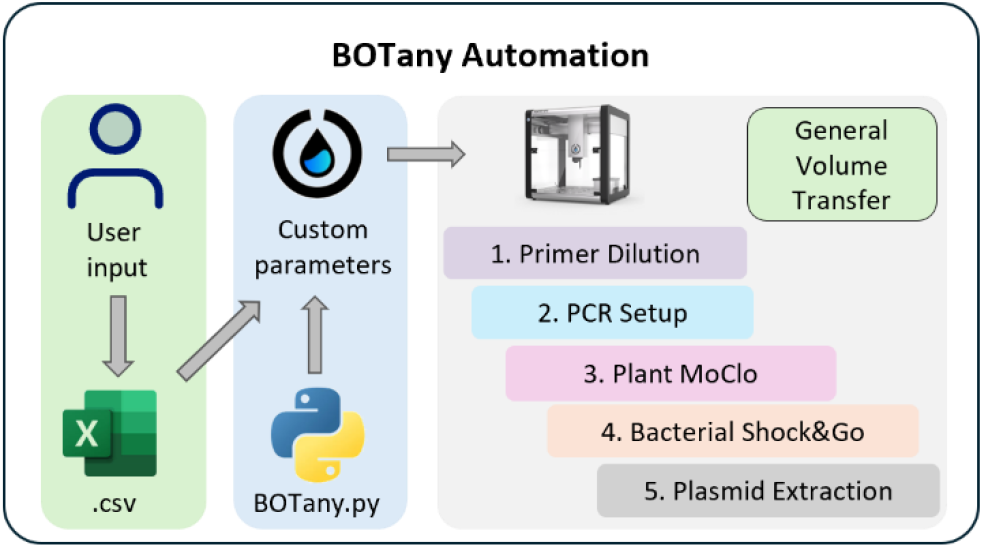

## Introduction

Despite the rise of generative artificial intelligence (AI), integration of robots in both industrial and domestic settings (2024), most students and bench scientists still carry out molecular biology techniques in an artisanal manner using manual pipettes. To clone a gene, assemble a multi-part construct, and/or transform an organism, success often depends on an individual’s prior experience and attention to detail. Pipetting liquids at the microliter (µL) scale is subject to technical errors, challenging to scale up, and can lead to repetitive stress injuries. Biofoundries can address these issues by offering a dedicated infrastructure for academic and industry stakeholders to automate common procedures in biology and bioengineering (Tellechea-Luzardo *et al*., 2022). A select group of institutions have set up a global alliance of biofoundries (Hillson *et al*., 2019) to speed up the design-build-test-learn (DBTL) cycle in synthetic biology. Biofoundries enable throughput experiments necessary to train AI models, accelerate scientific research, and empower product development, but are not available to most research-intensive universities in the United States and most other countries due to the high start-up and maintenance costs.

High-end laboratory robots (in excess of $100,000 per unit) typically require dedicated personnel, proprietary software with limited flexibility, expensive consumables, and/or mandatory service contracts (Torres-Acosta *et al*., 2022; Stephenson *et al*., 2023). To save costs, some labs have built do-it-yourself (DIY) systems that integrate entry-level robotic arms and 3D-printed components (Gome *et al*., 2019; Rupp *et al*., 2022). Because engineering and programming skills required to build and operate DIY robots pose barriers to biologists, modular benchtop liquid handlers such as the Opentrons OT-2 have become the go-to option for entry-level lab automation. The OT-2 robot consists of a modular chassis (resembling a biosafety cabinet) and 11 plug-and-play decks for standard labware plus add-on modules for sample cooling, shaking, magnet-based purification and thermocycling. As an example of their utility at scale, numerous OT-2 robots were used in research and clinical settings to automate SARS-CoV-2 diagnostic workflows and to test thousands of patient samples per day (Lázaro-Perona *et al*., 2021; Villanueva-Cañas *et al*., 2021). In the bioengineering community, several research groups have showcased the suitability and affordability of OT-2 robots for high-throughput DNA assembly (Storch *et al*., 2020; Bryant *et al*., 2023), *Agrobacterium* transformation (Annese *et al*., 2025), RNA extraction (Moufarrej and Quake, 2023), and single-cell proteomics (Montes *et al*., 2024), but comprehensive user-friendly solutions are still lacking for plant biologists.

While the open-source Opentrons application programming interface (API) supports Python-based protocol development and sharing, it still poses barriers to biologists and engineers without programming skills. Protocol Designer, an Opentrons web-based drag-and-drop interface, offers simple workflows for first-time or casual users, but it quickly becomes cumbersome to design complex liquid transfer schemes which require multiple changes in parameters and labware between experimental runs. From our hands-on experience and from personal communication with other synthetic biologists working with plants and microbes, many of the OT-2 protocols shared online (e.g. Opentrons Protocol Library) have not been experimentally validated and are not sufficiently accessible or versatile.

To tackle this hurdle, we developed a collection of OT-2 protocols that enable biologists, including undergraduate students interested in plant molecular biology, to automate several essential techniques by modifying spreadsheets. Instead of diving into the Python code or generating new scripts, users simply fill an Excel template with reagents, volumes, and transfer steps and save it as a Comma Separated Values (CSV) file that is imported to the Opentrons App. Our Python scripts parse the table and execute each instruction on the OT-2 robot, preserving full control over liquid handling without any user modifications to the code. By shifting protocol design into a familiar spreadsheet format, we offer a student-friendly interface that still delivers the flexibility and precision of custom scripting. The BOTany Methods automate primer dilution, PCR, DNA assembly, *E. coli* transformation, plasmid extraction, and more, thereby reducing hands-on time and increasing throughput for essential molecular biology tasks.

## Results

### CSV-based OT-2 protocols with runtime parameters

Our BOTany Methods can be applied to one or more robots, with separate units offering operational efficiency without changing the hardware between protocols (**Supplementary Fig. S1**). We initially built and/or modified custom OT-2 protocols in Python, but the manual editing of the code proved inefficient between experiments and challenging for lab members without training in computer science. To address this, protocol commands such as well locations and reagent definitions were migrated into Excel-editable CSV files. Our early implementation relied on manual copy-and-paste of CSV instructions into Python scripts, leaving room for errors.

Since the release of API v2.20 and App v7.3.0, Opentrons added the possibility to upload user-defined CSV files containing custom runtime parameters for Python scripts (**Table 1**). This shift to CSV-driven workflows enabled us to create more flexible protocols that require no coding knowledge or modification of the scripts, greatly shortening the learning curve to start automating molecular biology research and education for students and more experienced researchers.

**Table 1.** Summary of customizable parameters in BOTany protocols and Opentrons App. Protocol features: user-defined (✓), fixed in Python code (•); not available (-).

|  |  |  | BOTany1-Primers | BOTany2-PCR<br>BOTany3-MoClo | BOTany4-Shock&Go | Botany5-MagBead | BOTany6-Universal |
| --- | --- | --- | --- | --- | --- | --- | --- |
| BOTany<br>Methods<br>and CSV<br>Input | **Labware** | Module | • | • | • | • | ✓ |
|  |  | Labware | • | • | • | • | ✓ |
|  | **Reagents** | Location | • | ✓ | • | • | ✓ |
|  |  | Volume | ✓ | ✓ | ✓ | ✓ | ✓ |
|  | **Liquid**<br>**transfers** | Source | • | ✓ | ✓ | • | ✓ |
|  |  | Destination | • | ✓ | ✓ | • | ✓ |
|  |  | Transfer<br>volume | ✓ | ✓ | ✓ | • | ✓ |
|  |  | Tip use | • | ✓ | ✓ | • | ✓ |
| Opentrons<br>App<br>Parameters | **Pipettes** | Type | P300 | P20 | P20 and P300 | 8-channel<br>P300 | P20, P300,<br>P1000 |
|  |  | Mount | ✓ | ✓ | ✓ | ✓ | ✓ |
|  | **Starting tip location** |  | ✓ | ✓ | ✓ | ✓ | ✓ |
|  | **Thermocycler** |  | - | ✓ | - | - | - |
|  | **Sample number** |  | ✓ | - | - | ✓ | - |

### Automation of vector construction

#### From automated primer dilution to PCR

Synthetic DNA oligonucleotides (oligos or primers) serve as starting points for numerous molecular biology workflows including DNA synthesis, target sequence amplification for cloning or genotyping plants/microbes, and for Sanger sequencing. Since most labs order primers in a shelf-stable, lyophilized format that requires manual resuspension, we developed a fully automated protocol for the preparation of primer stocks (**Fig. 1A**). We use the BOTany1-Primers method as a practical solution that introduces new users to biological automation. First, the CSV file instructs the robot to add variable volumes of nuclease-free water to the lyophilized primer tubes whose final concentrations should be 100 µM. After pipette mixing the concentrated solutions, the robot also transfers aliquots to prepare 10 µM primer stocks in 1.5 mL tubes (**Supplementary Method 1**). We have successfully applied this method to process more than 200 primers for multiple users in our laboratory.

**Figure 1.**
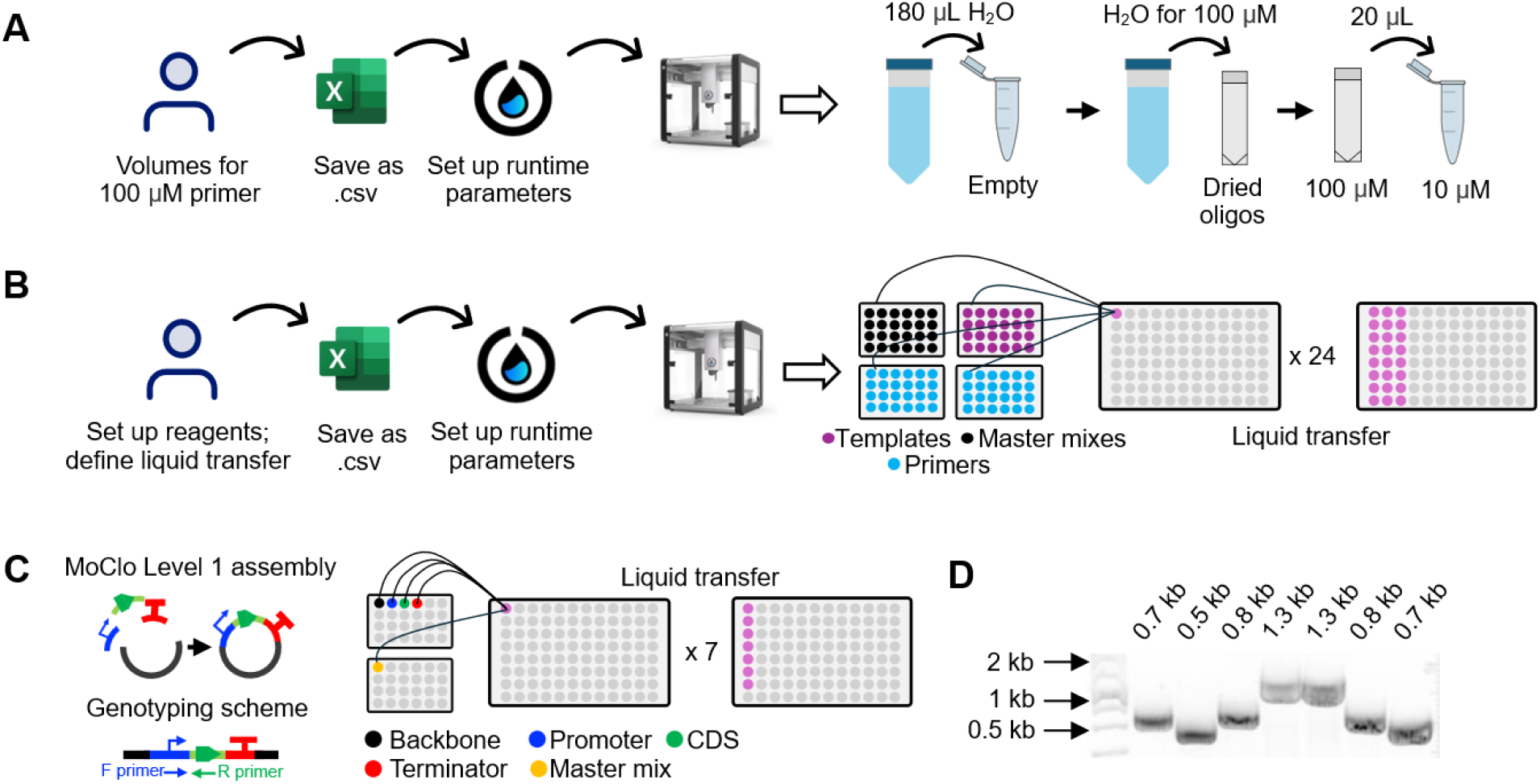
Workflow of molecular cloning. **A)** Automated primer dilution using BOTany1-Primers. **B)** DNA amplification using BOTany2-PCR methods. **C)** Plant Modular Cloning (MoClo) using BOTany3-MoClo methods, with MoClo Level 1 assemblies as an example. **D)** Colony PCR showing the expected amplicons for seven different Plant MoClo Level 1 assemblies from panel (C) using genotyping primers from (A).

Next, we developed BOTany2-PCR protocols (**Fig. 1B**) that support up to 120 reagent tubes in a 1.5 mL format (primers, templates, master mixes, etc.) to prepare reactions in 8-well PCR strips or in a 96-well plate. To accommodate different laboratory setups, there are two versions for the protocol to support the Opentrons thermocycler module in running on-deck reactions (version A) after all the liquid transfers are completed (**Supplementary Method 2**); version B supports off-deck thermocyclers from most other molecular biology companies. In both cases, a CSV file defines the pipetting instructions, including the liquids that will be loaded in the OT-2 and the desired transfer steps (e.g. source, destination, volume, tip and pipette choices). Sensitive reagents such as enzymes are chilled on-deck for the duration of the experiment using the on-deck temperature module. Although the OT-2 Temperature module can cool samples to 4°C, we have found it practical to pre-chill its aluminum blocks in the fridge and/or set the cooling temperature in the scripts to 14°C to strike a balance between time required to reach the setpoint (< 2 min in a room at 23°C) and the need to protect the reagents.

#### Automated Golden Gate assembly

Golden Gate assembly enables seamless, scarless assembly of multiple DNA fragments in a single reaction, making it ideal for constructing complex genetic constructs quickly and efficiently (Engler *et al*., 2008). Golden Gate standards have been established in the plant synthetic biology community (Patron *et al*., 2015), and collections such as the Plant Modular Cloning (MoClo) toolkit provide reusable coding sequences and regulatory elements for plants (Engler *et al*., 2014) and other hosts (Bird *et al*., 2022), including yeast (Püllmann *et al*., 2021; Grieß-Osowski *et al*., 2025). Plant MoClo automation improves throughput and accuracy, especially for early career researchers, parallel assembly of multiple constructs, or the generation of DNA libraries. Like the PCR protocols (**Fig. 1B**), we created fully automated BOTany3-MoClo versions for on-deck and off-deck thermocycling (**Fig. 1C**). In version A, users can specify the desired ligation time, temperature, and the number of digestion-ligation cycles for efficient Gate assembly. Seven Plant MoClo Level 1 vectors, consisting of a desired promoter, various coding sequences (cDNA), and a transcriptional terminator (**Fig. 1C**), were correctly assembled by the robot on the first try (**Fig. 1D**).

### Automated bacterial transformation

Automating the transformation of assembled plasmids into bacterial hosts such as *Escherichia coli* would increase the cloning success rate of undergraduate students and enable higher throughput projects. The BOTany4-Shock&Go protocol uses the Opentrons HEPA and Thermocycler modules (**Fig. 2A**) to perform up to 96 parallel *E. coli* transformations (**Supplementary Method S4**). First, the OT-2 robot aliquots competent cells (prepared in house) in the sterile PCR tubes, then adds newly assembled plasmid mixtures (**Fig. 2B**). The on-deck thermocycler heat shocks the cells, cools them, and finally aids their recovery by adding a nutrient-rich medium. The tubes are manually capped and then incubated on-deck for an hour before manual plating. To validate this protocol, we transformed eight Plant MoClo plasmid assemblies, ranging from 4 kb to 9 kb, and obtained around 30 to 160 colonies per plate using an aliquot of the recovered cells (**Fig. 2C**). Using blue/white colony screening, we found that fewer than 5% of colonies on each plate showed the *lacZ* activity of empty vectors (**Fig. 2D)**, except for one multimeric assembly with a low colony count. This protocol minimized the need for manual interventions (e.g. moving tubes between a water bath, an ice bucket, and a shaking incubator) and miniaturized the transformation volumes found in standard *E. coli* protocols. Our Shock&Go approach was also more efficient than the commercial Mix&Go! (Zymo Research) solution, which requires proprietary reagents for competent cell preparation but yielded fewer colony numbers without heat shock.

**Figure 2.**
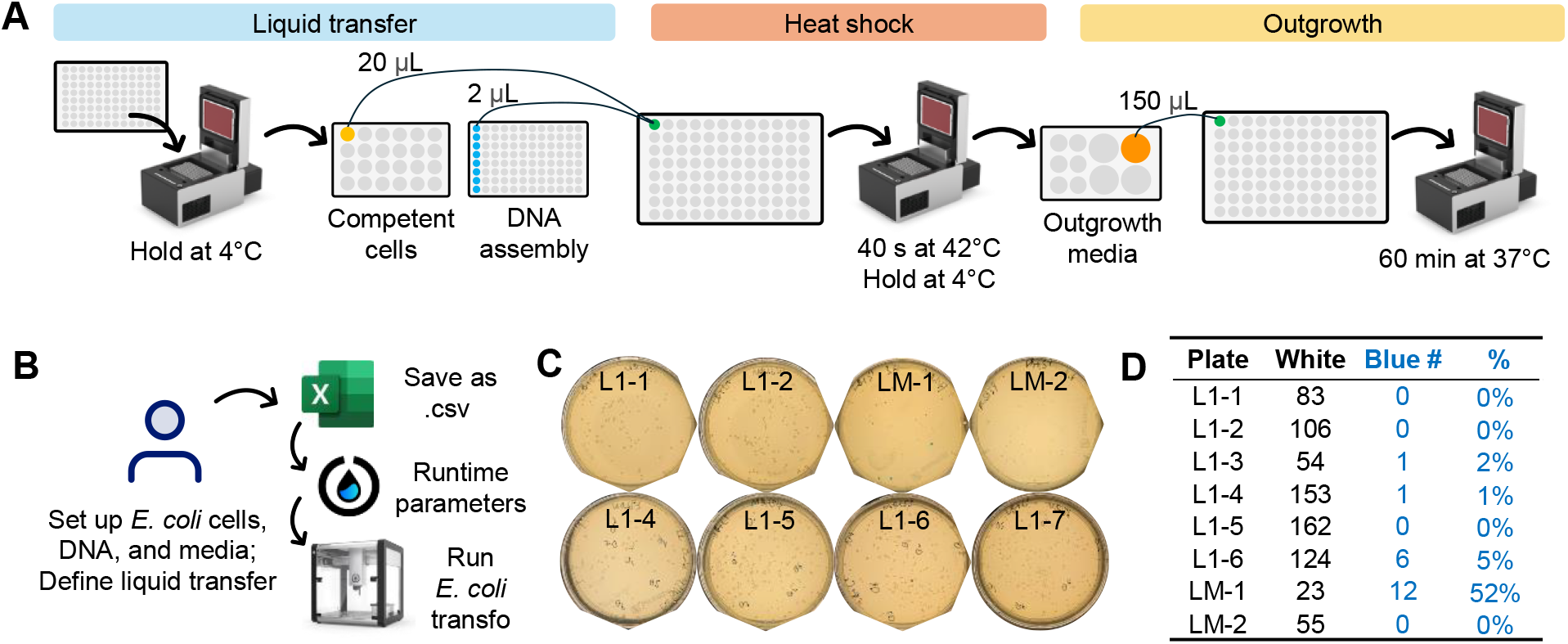
Automated *E. coli* transformation using BOTany4-Shock&Go. **A)** OT-2 workflow of the *E. coli* transformation protocol. **B)** Computer steps for the *E. coli* transformation protocol. **C)** Representative colony growth from 50 µL aliquots of transformed cells using different DNA assemblies. The L1 plates contain MoClo level 1 assemblies, while LM indicates MoClo level M assemblies that contain larger, multimeric constructs. **D)** Colony numbers after overnight growth and blue-white screening. The blue percentage denotes the empty vectors, and should be close to 0% for efficient assemblies.

### Semi-automated MagBead plasmid extraction

Most laboratories extract plasmid DNA using commercial miniprep kits containing spin columns, which require extensive pipetting and multiple centrifugation steps (plus capping and uncapping tubes), or the complete removal of solvent washes using a vacuum manifold. Our semi-automated protocol can process up to 96 samples using the Zyppy-96 Plasmid MagBead Miniprep kit, which offers cost parity on a per sample basis ($1.6 to $2.0 each) with traditional spin column kits. The kit uses two types of magnetic beads (MagBeads) to purify *E. coli* plasmid DNA and was initially selected based on the availability of a ‘compatible’ script in the Opentrons Protocol Library. However, neither the OT-2 scripts available online nor the standard kit instructions yielded sufficient DNA in our laboratory, so we had to develop a novel manual workflow and a Python script in consultation with Opentrons and Zymo. Neither the OT-2 Magnetic Module, nor the ZR-96 MagStand (recommended by Zymo) were strong enough to fully pellet the clearing MagBeads in 96-well plates. To address this, we pivoted to growing cells in autoclavable 24 deep-well plates that yield ample growth in standard laboratory shakers (e.g. 250 rpm at 1.9 cm orbit), are compatible with the Zymo MagStand, and provide extra space for initial pipetting steps that were difficult to automate (**Fig. 3A**). The BOTany5-MagBead method performs the remaining steps, including plasmid binding, washing, and elution using an 8-channel pipette (**Fig. 3B**). The Magnetic Module and the Heater Shaker are used to pull down the DNA during multiple washes and for incubations (**Supplementary Method 5**), respectively. Although the MagBead-extracted plasmids had only 5–10% of our typical miniprep (**Supplementary Fig. S2**), they displayed the correct restriction enzyme digestion patterns (**Supplementary Fig. S3**), along with some chromosomal DNA. We used MagBead-extracted plasmids (corresponding to coding sequences in Plant MoClo Level 0) for automated Golden Gate assembly and *E. coli* transformation using the aforementioned BOTany methods. PCR genotyping showed that all tested colonies contained the desired genes (**Fig. 3C**). Finally, we demonstrated that MagBead-extracted episomal plasmids (MoClo Level 1) were suitable for *Pichia* yeast cell electroporation (Lin-Cereghino *et al*., 2005), which requires more stringent DNA concentration and purity than bacterial transformation (**Fig. 3D**).

**Figure 3.**
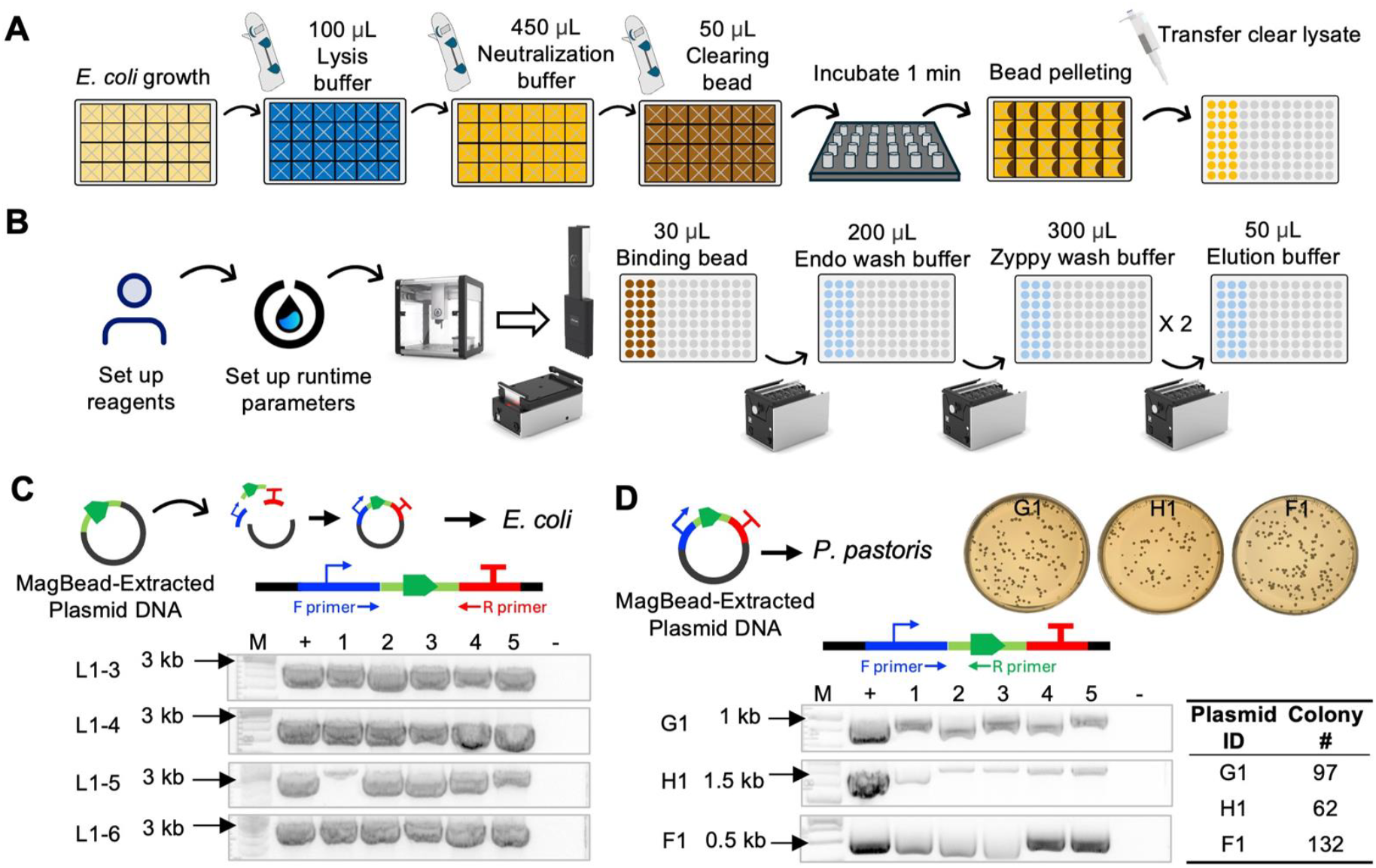
BOTany5-MagBead automation of plasmid extraction. **A)** Manual steps to obtain clear bacterial cell lysate. **B)** Workflow for automated binding, washing, and elution of plasmid. **C)** Golden Gate assembly using the MagBead-purified plasmids and *E. coli* transformation. **D)** *Pichia* electroporation using MagBead-extracted plasmids. Five colonies per transformation were genotyped in (C) and (D). ‘M’ indicates DNA markers, ‘-’ indicates negative control.

### A Versatile BOTany Protocol

In addition to the common procedures described above, we created a BOTany6-Universal pipetting protocol (**Fig. 4A**) that offers full flexibility in terms of hardware, labware, pipettes, tips, and liquid transfers (**Supplementary Method 6**). Our spreadsheet has drop-down selections with defined parts from the Opentrons API (e.g. tip rack, labware, and hardware modules) to simplify user input and avoid typing errors. This protocol was used to fill a 96 deep well plate with 500 µL of media and to inoculate wells with fluorescent *Pichia* yeast strains in a mosaic pattern that would be difficult to replicate by hand. After three days of cultivation, green and red fluorescent cells were only detected for the expected wells (**Fig. 4B**). We did not detect any cross-contamination or growth in nutrient-rich wells that were not inoculated with yeast cells, highlighting the efficiency of the HEPA module and accuracy of the pipetting steps.

**Figure 4.**
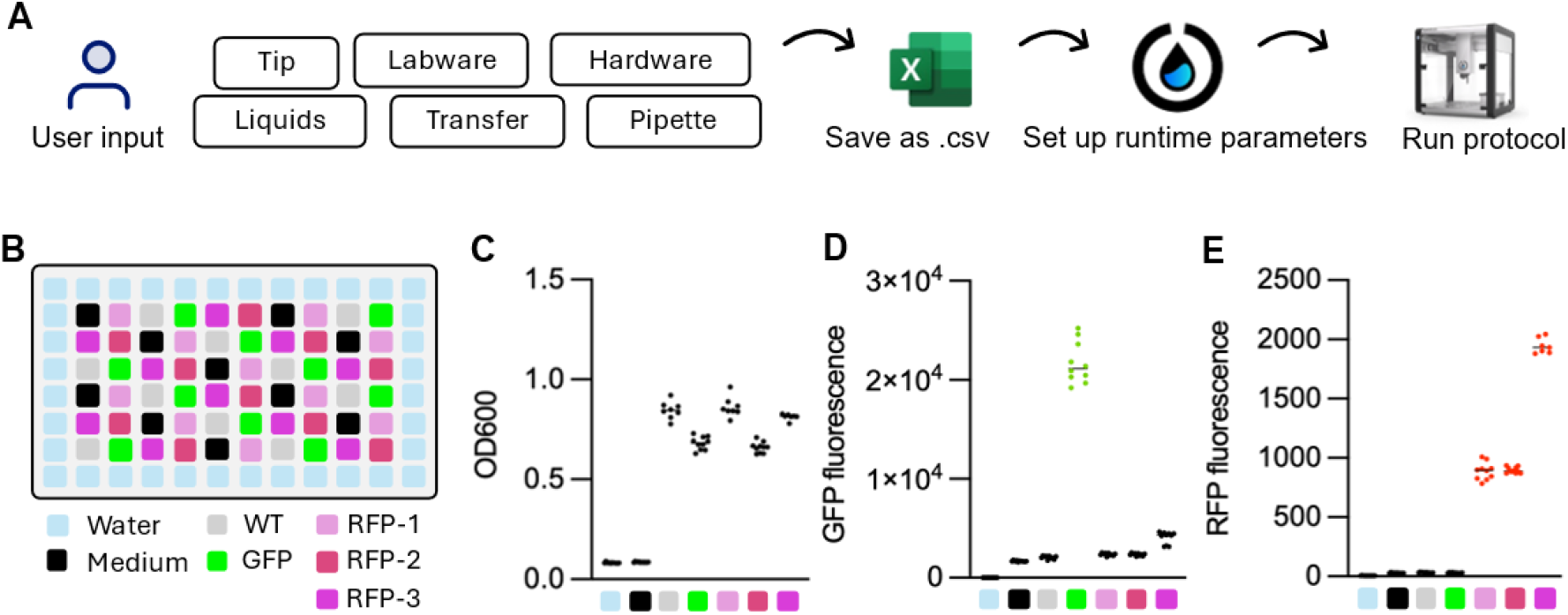
BOTany6-Universal liquid transfer. **A)** Workflow of a general liquid transfer protocol. **B)** Inoculation of a 96 deep well plate using different *Pichia* fluorescent cell strains and liquids in a mosaic pattern. After 72 h incubation at 800 rpm and 30°C, **C)** OD600, **D)** GFP fluorescence, and **E)** RFP fluorescence were measured using 1/10 diluted aliquots of yeast cell cultures.

## Discussion

Recently, the iBioFAB platform was programmed to automate tedious steps in plant transformation (Dong *et al*., 2025), such as protoplast preparation. However, high-end biofoundries remain inaccessible to most researchers because they are prohibitively expensive to replicate. Here, we present an affordable hardware setup and custom BOTany methods that automate several time-consuming steps in plant synthetic biology and can enable other labs to integrate liquid handlers in their research and teaching activities. The BOTany collection provides a streamlined approach and overcomes technical challenges encountered while teaching plant molecular biology methods to undergraduate students at the University of Florida. The Opentrons OT-2 system is currently one of the most affordable commercial solutions for liquid handling. The robot is sufficiently compact and robustly built to be occasionally transported, and we have successfully moved it on rolling carts for classroom demonstrations in neighboring labs or buildings. With considerable interest from industry, academic, and community science labs, full-time researchers as well as DYI biologists have promoted a protocol-sharing culture for the OT-2. Despite the convenience of the Opentrons Protocols Library (https://protocols.opentrons.com/), we caution that shared protocols should not be taken for granted since they may lack experimental validation or rigorous testing. For example, the Zymo MagBead protocol originally published by Opentrons failed for several reasons, such as inadequate Magnetic Module parameters. Peer-reviewed pipelines such as AssemblyTron (Bryant *et al*., 2023) are more robust, but require users to be proficient with multiple software and to modify scripts, which can deter plant biologists and other users.

Our implementation of customizable templates in Microsoft Excel overcomes programming hurdles and leverages software that is familiar to most students. The recently released OT-Mation also employs CSV files to generate Python protocols for routine OT-2 liquid transfer tasks that do not require advanced module control (Laverick *et al*., 2025). While this resembles the BOTany approach, OT-Mation did not provide detailed, biologically relevant examples. Meanwhile, our protocols offer a complete workflow from dry primers to the purification of newly assembled constructs. In essence, our platform enables the modular assembly of plant constructs (e.g. for single or multi-step enzymatic reactions), which can be quickly prototyped in heterologous hosts such as yeast before more time consuming experiments in model plants or crops (Grieß-Osowski *et al*., 2025). Users are only required to modify Excel CSV templates, since the Python code development and experimental validation was completed by our team. As a result, no additional software or scripts are needed: users simply load or drag our BOTany protocols and updated CSV files onto Opentrons App to execute the procedures, thereby providing a direct route to laboratory automation. We have rigorously validated each protocol step using concrete example workflows, demonstrating that each protocol works reliably under standard laboratory conditions. We also supply step-by-step instructions for each BOTany method, annotated screenshots and troubleshooting tips, so that researchers unfamiliar with the OT-2 interface can install and execute our protocols successfully from the first day.

In addition to OT-2 protocols dedicated to common procedures in synthetic biology, we also present a universal transfer protocol to serve as the foundation for additional tasks, such as RNA normalization and cDNA synthesis prior to real-time PCR. Our protocols and the OT-2 platform support a wide range of consumables, which provides experimental flexibility and does not restrict users to one vendor or proprietary supplies. Pipette tips from other manufacturers (e.g. universal 20 µL and 200 µL tips from Sarstedt) can be integrated seamlessly on the OT-2, and the system can accommodate custom labware files for plates or reservoirs. Although pipette tips and other modules must be calibrated during deck setup, at least during first-time use for a given protocol and slot, the OT-2 system bypasses the service contract required for more expensive robots, including the new Opentrons Flex. One benefit for more expensive robots such as the Flex is the option of a gripper to move labware between deck slots and modules. This could further automate our MagBead-based plasmid extraction method, which already yielded sufficient plasmids for DNA assembly and yeast transformation.

In conclusion, the BOTany methods combine affordable robots with modular, CSV-driven protocols to complete routine as well as advanced molecular biology tasks. Our detailed workflows are feasible to implement at other institutions and improve experimental throughput and reproducibility while remaining accessible to undergraduate students and other early-career researchers. Due to the modular nature of the OT-2 robots and the BOTany methods, the protocols are easy to get modified by the community and can be integrated with other recent methods for plant bioengineering (Annese *et al*., 2025). By lowering the barriers to start automating biology, our workflows for end-to-end molecular cloning will accelerate research and product development in plant sciences and beyond.

## Materials and methods

### BOTany method setup

The BOTany protocols were developed using Python framework of the Opentrons OT-2 API. Custom labware were measured using a digital caliper, and the.json files were created via link (https://labware.opentrons.com/#/create). **Supplementary Method 1** features a general guide for using the BOTany Methods and step-by-step instructions for the BOTany1-Primers protocol. **Supplementary Tables S1** and **S2** summarize the files and physical plasticware used in BOTany Methods. All the Python protocols, customizable tables for the CSV runtime parameters, and third-party labware files developed in this study can be found on our public GitHub repository (https://github.com/cvoiniciuc/BOTany).

### Primer dilution and PCR Setup

Synthetic oligos were synthesized by Eurofins Genomics (USA) in dry, salt-free, 25 nM scale, which typically require 100 to 300 µL water to prepare the 100mM stocks (**Supplementary Method S1**). The BOTany2-PCR-Table.xlsx template on GitHub illustrates 10 µL genotyping reactions containing 5 µL Red Taq 2X Master Mix (VWR International; 76620-474) and 2 µL water (together referred to as Master mix1), along with 1 µL of each 10 mM primer and 1 µL of DNA template. In this setup, Master mix1 is aliquoted first, followed by single transfers of primers and DNA templates. To save tips and time when genotyping many plant or microbial samples, primers could be pre-mixed with the master mix, leaving the DNA template as the only variable. This PCR setup can be scaled up to a 20 µL reaction volume and works with Phusion DNA Polymerase (Thermo Fisher Scientific) or other high-fidelity polymerases for cloning.

### Golden Gate assembly

DNA parts provided in the MoClo Toolkit (Addgene kit # 1000000044) were a gift from Sylvestre Marillonnet. Our Golden Gate reactions used FastDigest Eco31I (BsaI), and BpiI (BbsI) type IIS restriction enzymes (Thermo Fisher Scientific), but the protocols are compatible with alternative suppliers (New England Biolabs) or enzymes. Efficient ligation was obtained with T4 DNA ligases from Thermo Fisher Scientific (MA, USA; 5 U/µL; EL0012) at 22°C, and New England Biolabs (MA, USA; M0202) at 16°C. Both suppliers provided efficient assembly in our methods.

In BOTany3-MoClo-Table.xlsx, the example illustrates 10 µL Golden Gate reactions consisting of 1 µL each of the MoClo parts (backbone, promoter, CDS, and terminator), combined with 6 µL of Master Mix containing 1 µL 10 mM ATP (Thermo Fisher Scientific, MA, USA; R0441), 1 µL 10× FastDigest Buffer, 0.5 µL FastDigest type IIS restriction enzyme, 0.5 µL T4 DNA ligase (either brand mentioned above), and 3 µL water.

### *E. coli* Methods

To prepare competent cells, 10 mL overnight culture from a single TOP10F’ colony was used to inoculate 1 L of low salt lysogeny broth (LB) medium. E. coli was grown at 37°C and 250 rpm in an Innova 42 shaker (Eppendorf) for 3 to 4 h until OD_600_ reached 0.4 and then chilled on ice for 30 min. All further steps were performed on ice or using equipment and reagents chilled to 4°C. Cells were divided into four 250 mL aliquots in pre-chilled centrifuge bottles and pelleted at 2,000 *g* for 15 min at 4°C. Each pellet was resuspended in 100 mL of 0.1 M MgCl_2_, and suspensions were combined into two bottles. Cells were pelleted again by centrifuging (as above) and resuspended in 100 mL ice-cold 0.1 M CaCl_2_. After incubation on ice for ≥20 min, cells were pelleted again and resuspended in 25 mL ice-cold 85 mM CaCl_2_ with 15% (v/v) glycerol. The two suspensions were combined, pelleted once more (1,000 × g, 15 min, 4 °C), and finally resuspended in 2 mL ice-cold 85 mM CaCl_2_ with 15% (v/v) glycerol to a final OD_600_ of 200 to 250. Aliquots (50 to 100 µL) were dispensed into pre-chilled sterile 1.5-mL tubes, snap-frozen in liquid nitrogen, and stored at −80°C.

The transformation efficiency of these cells was approximately 10^6^ colonies per µg of pUC19 plasmid DNA. After outgrowth on OT-2, 50 µL *E. coli* cell cultures were plated on 60 mm plates containing low salt LB agar supplemented with proper antibiotics (50 µg/mL carbenicillin and spectinomycin for MoClo L1 and LM constructs, respectively). For blue/white screening, 10 µL of X-gal stock (20 mg/mL in dimethyl sulfoxide) and 30 µL of IPTG stock (Isopropyl β-D-1-thiogalactopyranoside; 100 mM in water) were spread onto each 60-mm LB agar plate together with the transformed *E. coli* culture. Plates were incubated overnight at 37°C to allow colony formation.

### MagBead plasmid extraction

Positively genotyped *E. coli* colonies were inoculated in 1.6 mL low-salt LB medium with antibiotics in 24 deep-well plate sealed with air-permeable cover (Sigma Aldrich). The cultures were incubated overnight at 250 rpm at 37°C in an Innova® 42 shaker. For each culture, 800 µL was mixed with 200 µL of 75% glycerol to prepare glycerol stocks, which were stored in 1.6-mL CryoPure tubes (Sarstedt), and the rest of the culture was used for Zyppy-96 Plasmid MagBead Miniprep kit (Zymo Research, Cat# D4102) in **Supplementary Method S5**. After the automated steps, beads may be present in the elution plate but did not interfere with plasmid use, including the DNA quantification using a NanoDrop spectrophotometer.

In the traditional spin-column miniprep approach, miniprep spin columns (Syd Labs, MA, USA; MB082-PC200) were used in combination with the Qiagen Plasmid Miniprep Buffer Set (Qiagen, Germany; 27104). The same column was also compatible with buffers prepared in-house.

### Yeast growth for universal method validation

*Pichia* experiments were conducted as previously described (Grieß-Osowski et al., 2025), using strains expressing sfGFP and mScarlet fluorescent reporters under the methanol-inducible *pAOX1* promoter. Transformed GFP, RFP-1, RFP-2, RFP-3 lines along with X-33 strain were grown in 500 µL YP media supplemented with 1.5% (v/v) methanol, 0.5% (v/v) glycerol and 150 µg/mL hygromycin in a NEST 96-well 2 mL plate using a Multitron shaker (Infors HT, Switzerland) at 30°C, 800 rpm, 80% humidity for 72 h. Optical density and fluorescent signals were quantified on a BioTek Synergy H1 Microplate Reader (Grieß-Osowski *et al*., 2025).

## Acknowledgements

The initial BOTany setup was supported by an infrastructure grant from the University of Florida Institute of Food and Agricultural Sciences (UF/IFAS) and matching personnel funds from the Horticultural Sciences Department, which were secured together with Andrew D. Hanson. A.L. and L.L. were also supported by the UF AI Scholars and the UF/IFAS Summer Research Internships programs, respectively. We also acknowledge the Opentrons Education Initiative for helping us bring automation to the classroom through the donation of a teaching robot and support with the MagBead Python script to enhance the plant molecular biology curriculum.

## Author contributions

M.Q. led the implementation of the protocols, the manuscript drafting, the data acquisition and analysis. A.L. programmed the Python scripts and runtime parameters, and L.L. performed biological experiments using the robots. C.V. conceived the project, supervised and supported the method development, analyzed data, and revised the manuscript.

## Data availability

Data is provided in the supplementary section, with additional Python scripts, tables and custom labware files publicly available on GitHub (https://github.com/cvoiniciuc/BOTany). Additional information is available upon request from the corresponding author.

## Supplementary data

The following materials are available in the online version of this article.

**Supplementary Table S1**. Summary of BOTany Excel and Python files on GitHub.

**Supplementary Table S2**. Plasticware used in BOTany methods.

**Supplementary Table S3**. Yield and purity of MagBead-extracted plasmids.

**Supplementary Figure S1**. BOTany Foundry Configuration.

**Supplementary Figure S2**. Plasmids extracted with MagBead versus spin columns.

**Supplementary Figure S3**. Test digestions of MagBead-extracted plasmids.

**Supplementary Method 1**. General Guide and BOTany1-Primers Instructions.

**Supplementary Method 2**. BOTany2-PCR Instructions.

**Supplementary Method 3**. BOTany3-MoClo Instructions.

**Supplementary Method 4**. BOTany4-Shock&Go Instructions.

**Supplementary Method 5**. BOTany5-MagBead Instructions.

**Supplementary Method 6**. BOTany6-Universal Instructions.

## References

Annese D, Romani F, Grandellis C, Ives L, Frangedakis E, Buson FX, Molloy JC, Haseloff J (2025) Semi-automated workflow for high-throughput Agrobacterium-mediated plant transformation. Plant J Cell Mol Biol 122: e70118

Bird JE, Marles-Wright J, Giachino A (2022) A User’s Guide to Golden Gate Cloning Methods and Standards. ACS Synth Biol acssynbio.2c00355

Bryant JA Jr, Kellinger M, Longmire C, Miller R, Wright RC (2023) AssemblyTron: flexible automation of DNA assembly with Opentrons OT-2 lab robots. Synth Biol 8: ysac032

Dong J, Croslow SW, Lane ST, Castro DC, Blanford J, Zhou S, Park K, Burgess S, Root M, Cahoon EB, et al (2025) Enhancing lipid production in plant cells through automated high-throughput genome engineering and phenotyping. Plant Cell 37: koaf026

Engler C, Kandzia R, Marillonnet S (2008) A One Pot, One Step, Precision Cloning Method with High Throughput Capability. PLoS ONE 3: e3647

Engler C, Youles M, Gruetzner R, Ehnert T-M, Werner S, Jones JDG, Patron NJ, Marillonnet S (2014) A Golden Gate Modular Cloning Toolbox for Plants. ACS Synth Biol 3: 839–843

Gome G, Waksberg J, Grishko A, Wald IY, Zuckerman O (2019) OpenLH: Open Liquid-Handling System for Creative Experimentation with Biology. Proc. Thirteen. Int. Conf. Tangible Embed. Embodied Interact. Association for Computing Machinery, New York, NY, USA, pp 55–64

Grieß-Osowski A, Robert M, Qiande M, Clauss S, Voiniciuc C (2025) Illuminating Glucomannan Synthases To Explore Cell Wall Synthesis Bottlenecks. ACS Synth Biol 14: 2472–2479

Hillson N, Caddick M, Cai Y, Carrasco JA, Chang MW, Curach NC, Bell DJ, Le Feuvre R, Friedman DC, Fu X, et al (2019) Building a global alliance of biofoundries. Nat Commun 10: 2040

Laverick A, Convey K, Harrison C, Tomlinson J, Stach J, Howard TP (2025) OT-Mation: an open-source code for parsing CSV files into Python scripts for control of OT-2 liquid-handling robotics. Synth Biol 10: ysaf009

Lázaro-Perona F, Rodriguez-Antolín C, Alguacil-Guillén M, Gutiérrez-Arroyo A, Mingorance J, García-Rodriguez J, Group on behalf of the S-C-2 W (2021) Evaluation of two automated low-cost RNA extraction protocols for SARS-CoV-2 detection. PLOS ONE 16: e0246302

Lin-Cereghino J, Wong WW, Xiong S, Giang W, Luong LT, Vu J, Johnson SD, Lin-Cereghino GP (2005) Condensed protocol for competent cell preparation and transformation of the methylotrophic yeast Pichia pastoris. BioTechniques 38: 44–48

Montes C, Zhang J, Nolan TM, Walley JW (2024) Single-cell proteomics differentiates Arabidopsis root cell types. New Phytol 244: 1750–1759

Moufarrej MN, Quake SR (2023) An inexpensive semi-automated sample processing pipeline for cell-free RNA extraction. Nat Protoc 18: 2772–2793

Patron NJ, Orzaez D, Marillonnet S, Warzecha H, Matthewman C, Youles M, Raitskin O, Leveau A, Farré G, Rogers C, et al (2015) Standards for plant synthetic biology: a common syntax for exchange of DNA parts. New Phytol 208: 13–19

Püllmann P, Knorrscheidt A, Münch J, Palme PR, Hoehenwarter W, Marillonnet S, Alcalde M, Westermann B, Weissenborn MJ (2021) A modular two yeast species secretion system for the production and preparative application of unspecific peroxygenases. Commun Biol 4: 1–20

Rupp N, Peschke K, Köppl M, Drissner D, Zuchner T (2022) Establishment of low-cost laboratory automation processes using AutoIt and 4-axis robots. SLAS Technol 27: 312–318

Stephenson A, Lastra L, Nguyen B, Chen Y-J, Nivala J, Ceze L, Strauss K (2023) Physical laboratory automation in synthetic biology. ACS Synth Biol 12: 3156–3169

Storch M, Haines MC, Baldwin GS (2020) DNA-BOT: a low-cost, automated DNA assembly platform for synthetic biology. Synth Biol 5: ysaa010

Tellechea-Luzardo J, Otero-Muras I, Goñi-Moreno A, Carbonell P (2022) Fast biofoundries: coping with the challenges of biomanufacturing. Trends Biotechnol 40: 831–842

Torres-Acosta MA, Lye GJ, Dikicioglu D (2022) Automated liquid-handling operations for robust, resilient, and efficient bio-based laboratory practices. Biochem Eng J 188: 108713

Villanueva-Cañas JL, Gonzalez-Roca E, Gastaminza Unanue A, Titos E, Martínez Yoldi MJ, Vergara Gómez A, Puig-Butillé JA (2021) Implementation of an open-source robotic platform for SARS-CoV-2 testing by real-time RT-PCR. PloS One 16: e0252509

