## Supplementary material for "BOTany Methods: Accessible Automation for Plant Synthetic Biology": Supplementary Method S1.pdf

---

### General Guide to BOTany Methods

##### Opentrons App

1. Opentrons provides a free app which is used to run the OT-2. It can be downloaded here: Opentrons App Download.
2. Note that the "API" for anything is from Opentrons. They can be found in the Opentrons Labware Library.

##### Starting Tip Selection

This feature can be useful if your tip rack is not full, and you do not want to manually replace the tips starting from A1.

1. If you use a new tip box, the starting tip location is A1 by default. The tips will be taken in an order of A1-B1-...H1-A2-B2...H2-A3, etc. If you have used the first 25 tips, the starting position would be B4, which can be specified in the runtime parameters.
2. Although our protocols allow the user to choose the starting tip position, a tip is required on A1 position if you plan to calibrate the OT-2's offsets. This is because the Opentrons only uses the A1 tip for labware calibration, not the user-defined starting tip.

##### Sample Placement Order

This only applies to the ***BOTany1-Primers*** method, where samples should be placed in an order of A1-A2-...A6-B1-B2-...B6-C1..., etc. In other words, row by row, top down.

##### Verification Steps on Opentrons App

After configuring the runtime parameters, there are four main setup steps on Opentrons App. Once these are completed, the robot will be ready to begin.

Screenshot of software page after runtime parameter setup

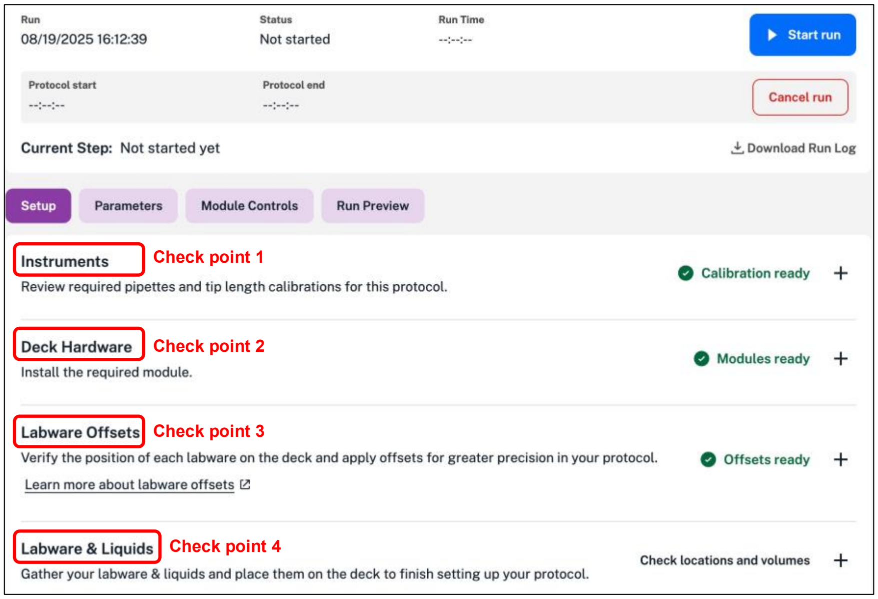

1. In **Instruments**, calibrate the deck and pipette. This is robot-dependent, therefore it is mandatory for initial robot set-up. If this data is not available, the Opentrons app will provide prompts to set this up. For future runs on the same robot, no more calibration is needed.

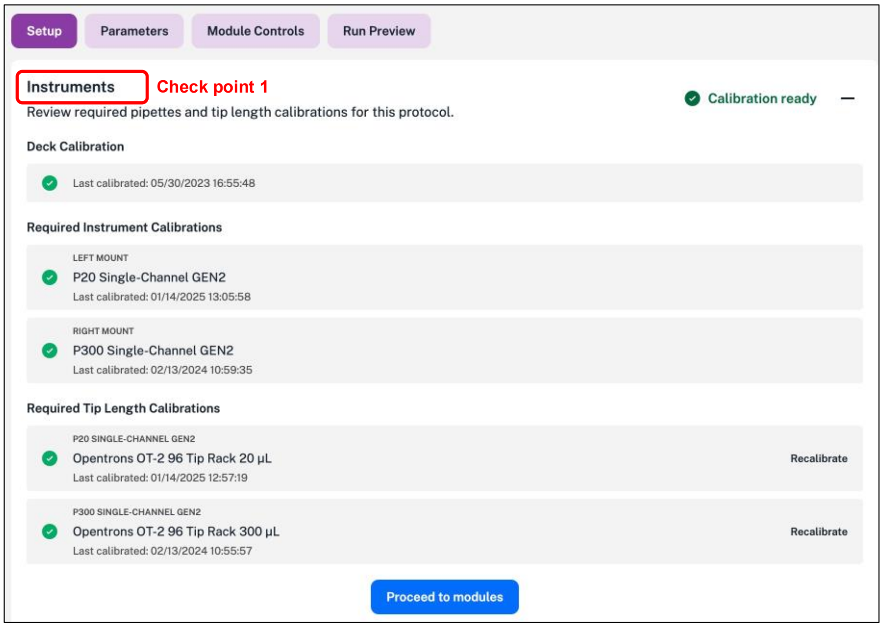

2. In **Deck Hardware**, place the required hardware on correct deck positions.

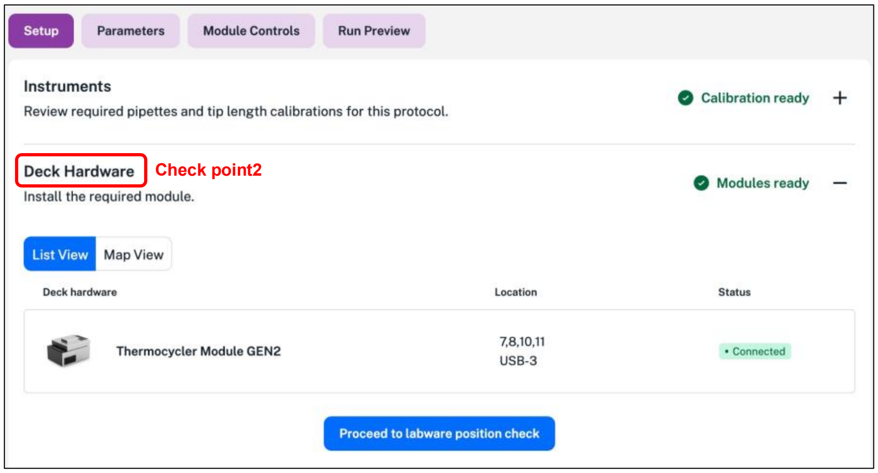

3. In **Labware Offsets**, run Labware Position Check to ensure the pipettes interact properly with each labware on the deck. It is mandatory for the initial run and protocol-dependent, remaining valid for up to 20 subsequent runs. Users can follow prompts built in the Opentrons App or this detailed online walk-through: Creating labware offsets with Labware Position Check.

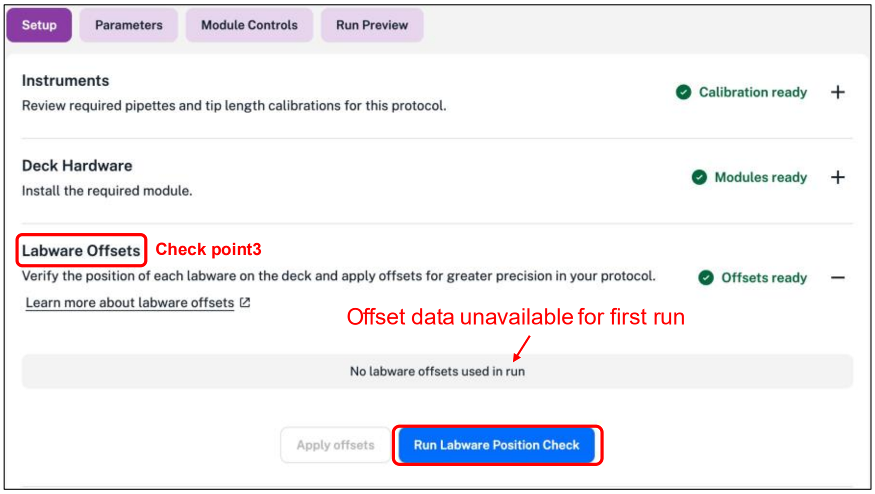

Before labware position check

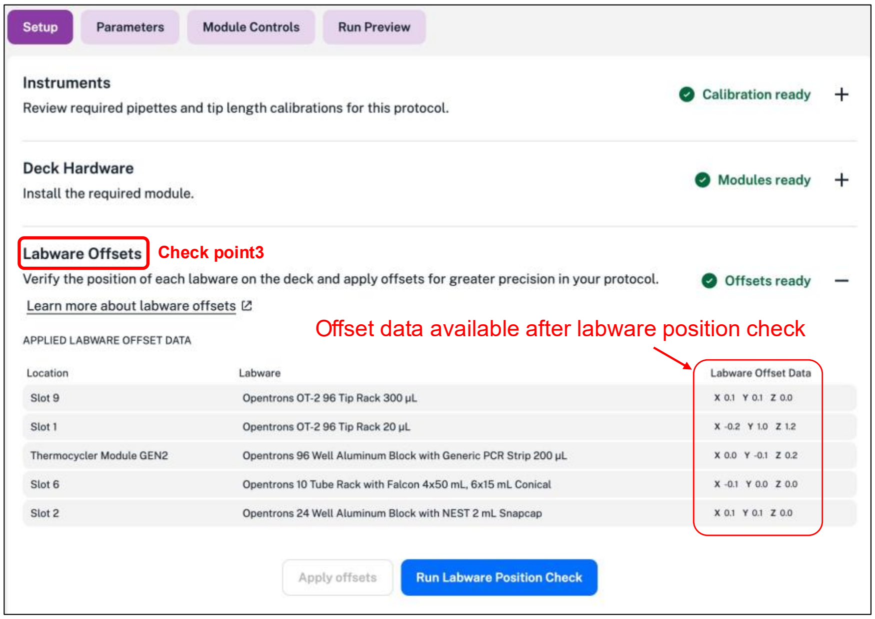

After labware position check

4. In **Labware & Liquid**, a graphical overview of the labware and liquids allows the user to load and check that everything is in the correct position.

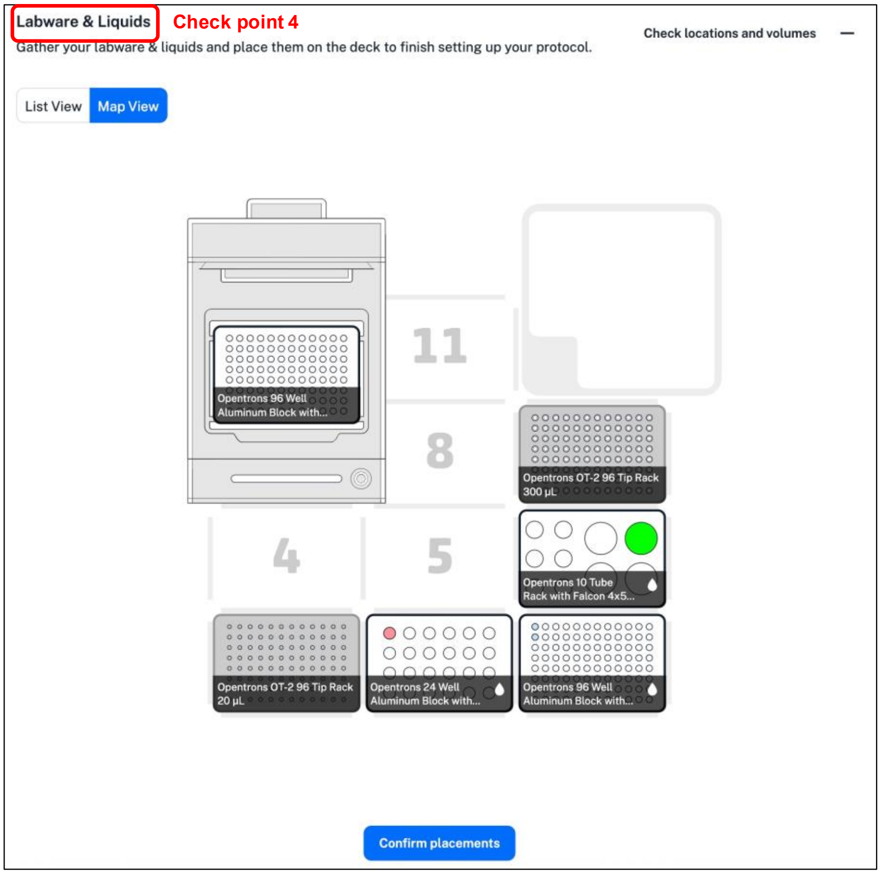

**Troubleshooting**

1. A calibration error will occur if the wrong pipette is chosen in the parameter setup.
   a. If this happens, cancel the run and start again with selection of the runtime parameters. Select the correct pipette(s) that matches your .CSV file and protocol.
2. An error like “Error: none type is not scriptable” most likely indicates an issue where a mismatch exists between the labware names (or syntax) defined in either the code/labware definition tables, and the liquid transfer table.
3. Sometimes a protocol may take a minute or two to process in the Opentrons app, which is normal for more complex protocols.
4. If a computer has problems connecting to the OT-2, one method to troubleshoot is to simply turn the OT-2 off and then back on. This usually resolves any connection errors.
5. If a protocol is loaded on the robot and the setup is in progress, the run must be cancelled (Cancel run button in the screenshot below) before the robot is available to load a new method. As described in each detailed method, the robot checks if any hardware is missing.

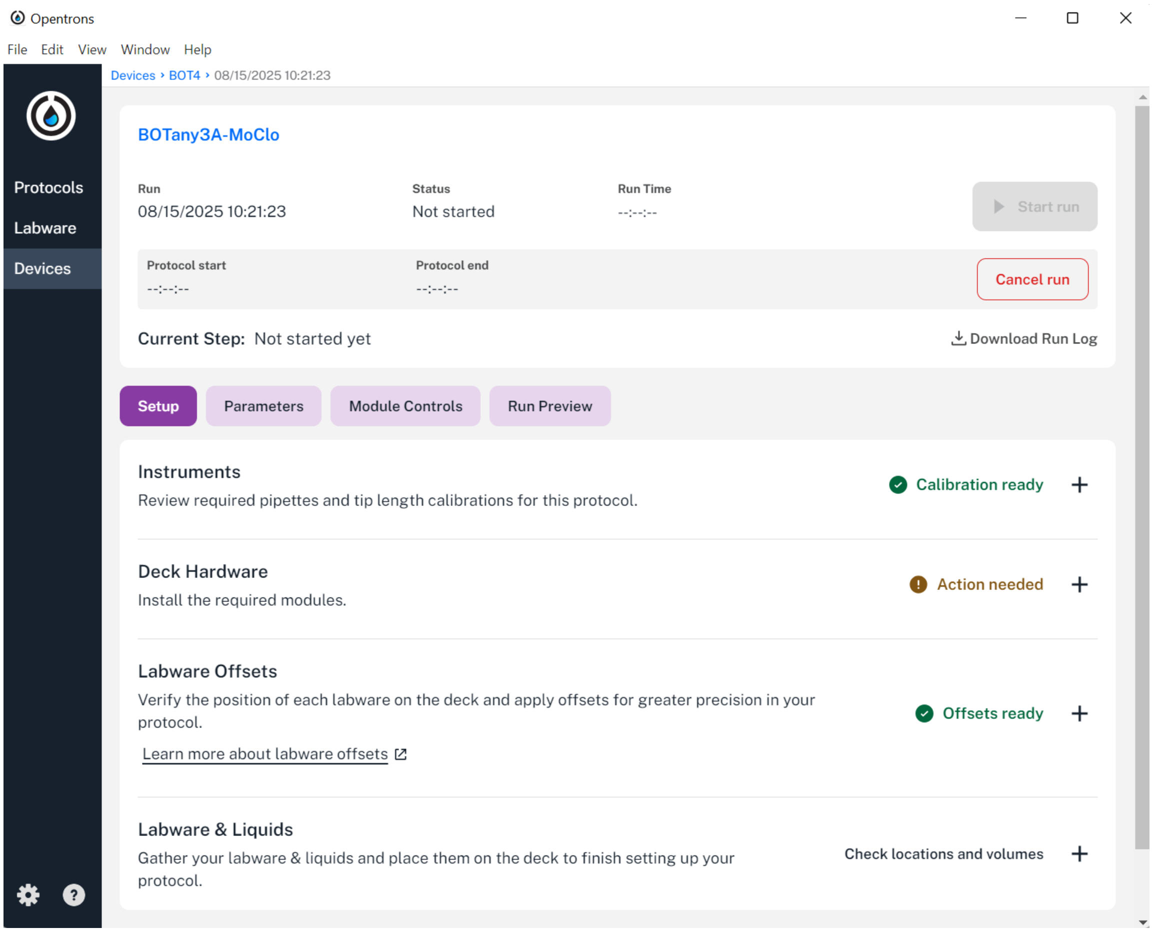

6. Except for the **BOTany5-MagBead** protocol, our methods do not currently use multichannel pipettes. The protocols allow you to select one in the runtime parameters if it is already mounted (but unused) on the OT-2.
   The OT-2 Protocol Designer can be used for simple experiments using 8-channel pipettes.

### BOTany1-Primers

This protocol fully automates the resuspension and dilution of up to 24 dry primers.

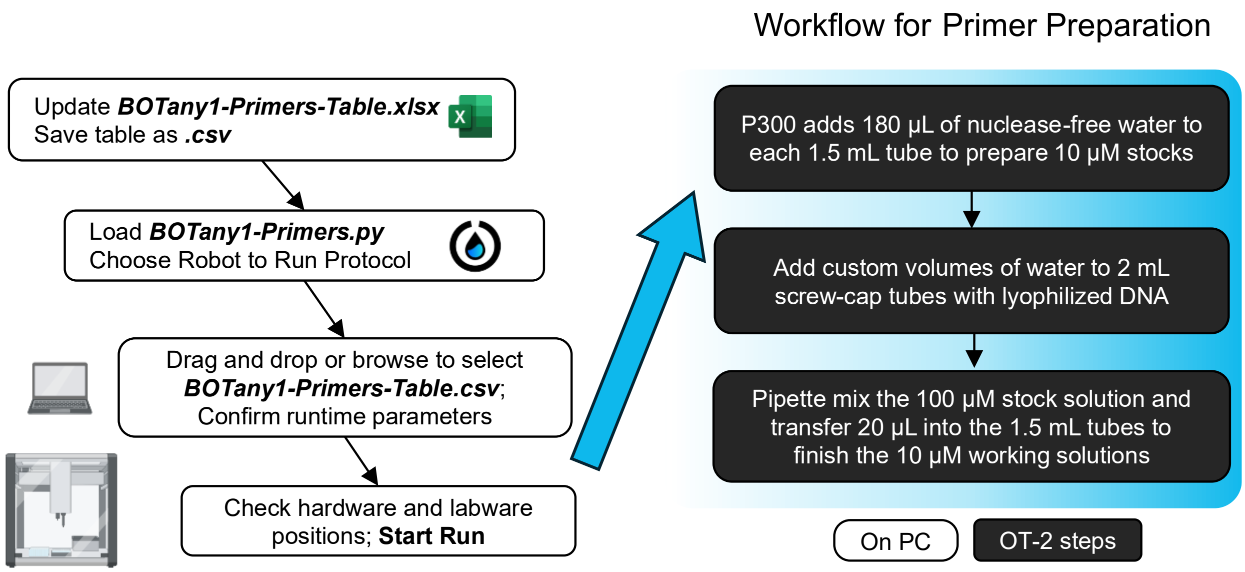

| **Required Hardware** | **API Name** |
|---|---|
| ○ P300 pipette | p300_single_gen2 |

| **Labware Quantity** | **API Name** |
|---|---|
| ○ 1 | opentrons_10_tuberack_falcon_4x50ml_6x15ml_conical |
| ○ 1 | opentrons_24_tuberack_nest_1.5ml_snapcap |
| ○ 1 | opentrons_24_tuberack_generic_2ml_screwcap |

##### BOTany1-Primers Instructions

1. Open the ***BOTany1-Primers-Table.xlsx*** template in Excel.

2. Edit **Pipetting Steps** (blue headings) to define the number of samples.

| | |
|---|---|
| **Source_Labware** | case-sensitive name of the labware to aspirate water from |
| **Source_Well** | position of well/tube within the Source_Labware |
| **Destination_Labware** | exact name of labware with screw-cap tubes (dry primers) |
| **Destination_Well** | destination well/tube within the Destination_Labware |
| **Transfer_Volume** | liquid transfer volumes in microliters (µL) |
| **Pick_Up_Tip** | TRUE = new tips, FALSE = re-use tip |

3. Save the updated Excel table in the .CSV format file [CSV UTF-8 format].
4. Open the Opentrons app. Import the ***BOTany1-Primers.py*** script in the Protocols tab.
5. Click the three vertical dots on the top right of the protocol to **Start setup**.
   a. Choose Robot to set up runtime parameters (select from available wireless or wired connections; to become available, OT-2 robot must complete/cancel prior methods).
   b. Drag and drop or browse for the .CSV file with liquid definitions and transfer steps.

c. Check or modify the remaining runtime parameters.

**Starting Tip Letter** and **Start Tip Number**: Location of the first row and column for the pipette tip box; The pipette picks up new tips by travel down each column, then shifting to the right. This feature is helpful if your tip rack is not full. If full, start with position A1.
**Pipette Location**: Choosing the Left/Right location of the required P300 pipette on the robot, from the perspective of someone standing in front of it.
**Other Pipette Choice** and **Other Pipette Location**: Select “None” or specify which additional pipette is already mounted and its location. The robot will not use the other pipette during this method.

6. After confirming the runtime parameters, verify the final robot setup (refer to *General Guide to BOTany Methods: Verification Steps on Opentrons App*). At this stage, the deck should be arranged as shown below.

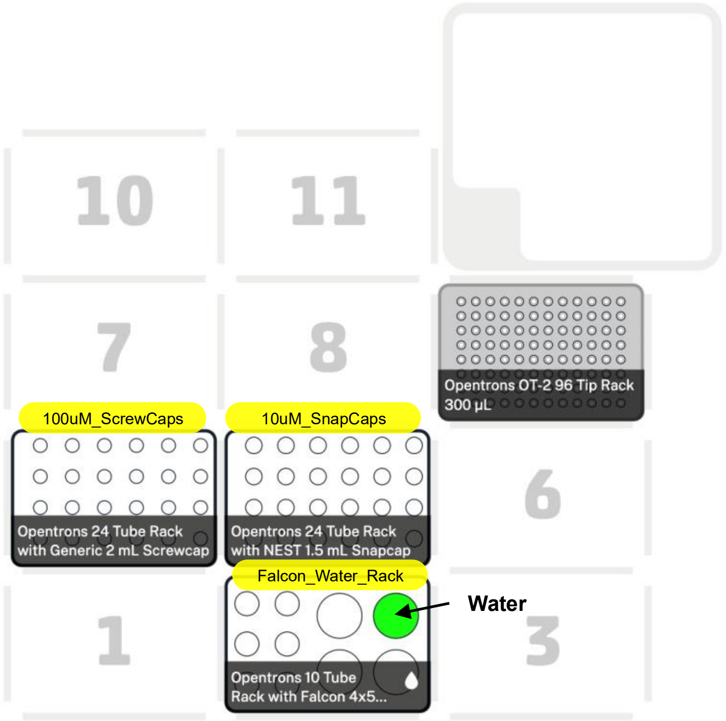

7. After checking the verification steps, click **Start Run** and enjoy the automated protocol.
