## Supplementary material for "BOTany Methods: Accessible Automation for Plant Synthetic Biology": Supplementary Method S2.pdf

---

#### BOTany2A-PCR for OT-2 Thermocycler

This protocol fully automates PCR reactions in a 96-well plate using up to 120 DNA components and enzyme mixtures loaded in 1.5 mL tubes. After pipetting the user-defined parts in the desired order, the robot will automatically seal the plate lid and run the PCR using the on-deck Opentrons Thermocycler.

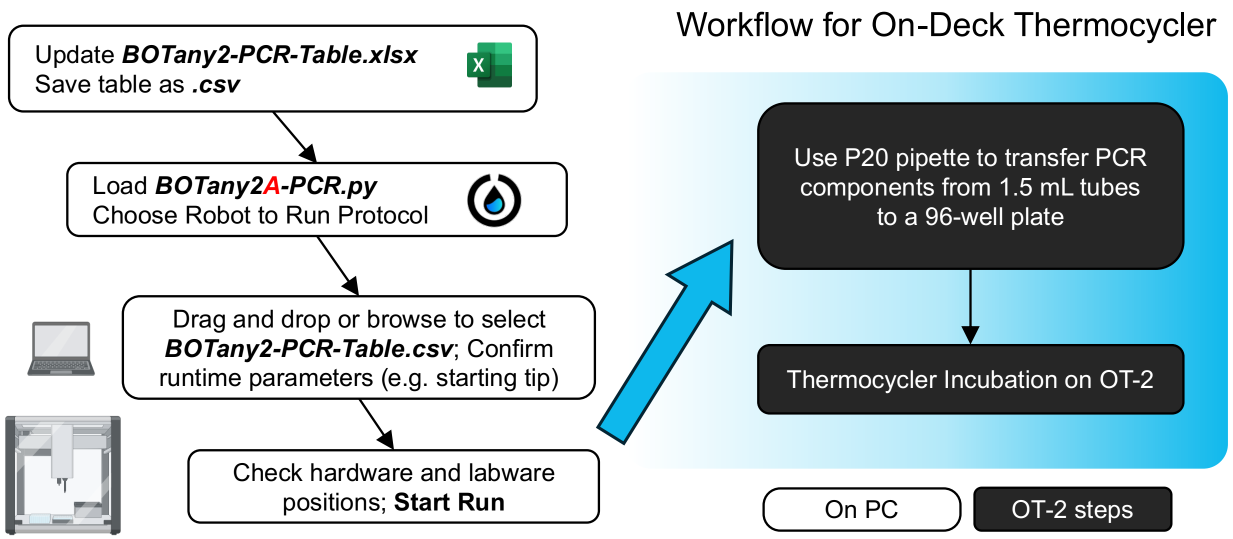

| **Required Hardware** | **API Name** |
|---|---|
| ○ P20 pipette | p20_single_gen2 |
| ○ Cooling block | temperature module gen2 |
| ○ Thermocycler | thermocyclerModuleV2 |

| **Labware Quantity** | **API Name** |
|---|---|
| ○ 1, pre-chilled[1] | opentrons_24_aluminumblock_nest_1.5ml_snapcap |
| ○ 1 to 4 | opentrons_24_tuberack_nest_1.5ml_snapcap |
| ○ 1 | nest_96_wellplate_100ul_pcr_full_skirt |

---

[1] Pre-chilling aluminum blocks in a fridge or on ice can expedite the cooling of temperature module.

#### BOTany2A-PCR Instructions

1. Open the ***BOTany2-PCR-Table.xlsx*** template in Excel.
2. PCR parts are stored in 1.5 mL snap-cap tube racks (labelled tube_rack1 to tube_rack4). Each rack can hold up to 24 tubes (for up to 96 input parts). The temperature module (labelled temp_tubes) can chill up to 24 additional tubes at 14°C.
3. Edit the **Initial Liquid Definitions** (green headings) to specify which liquids will be loaded in the robot. *Note:* OT-2 cannot sense if liquids are loaded correctly or their volumes.

| | |
|---|---|
| **Labware** | case-sensitive names used within the code; use drop-down menu |
| **Initial_Wells** | location of the liquid well/tube within a particular labware |
| **Initial_Volume** | the initial volume of said liquid in µL |
| **Liquid_Name** | any name you wish to assign the liquid |
| **Description** | any description you wish to assign the liquid (optional) |
| **Color** | HEX color codes that will show up in the OT-2 software deck view |

**Critical point:** Use the drop-down for each labware in Excel or copy-paste desired part names to ensure the correct syntax is used.

4. Edit **Pipetting Steps** (blue headings) to define volume transfers and robotic movements

| | |
|---|---|
| **Source_Labware** | case-sensitive name of the labware to aspirate (take) liquid from |
| **Source_Well** | position of well/tube within the Source_Labware |
| **Destination_Labware** | exact name of labware to dispense liquid into |
| **Destination_Well** | destination well/tube within the Destination_Labware |
| **Transfer_Volume** | liquid transfer volumes in microliters (µL) |
| **Pick_Up_Tip** | TRUE = new tips, FALSE = re-use tip (contamination risk) |
| **Pipette_Choice** | Left/Right location of required pipette (when facing the OT-2 front) |

5. Save the updated Excel table in the .CSV format file [CSV UTF-8 format].
6. Open the Opentrons app. Import the ***BOTany2A-PCR.py*** script in the Protocols tab.

7. Click the three vertical dots on the top right of the protocol to **Start setup**.
   a. Choose Robot to set up runtime parameters (select from available wireless or wired connections; to become available, OT-2 robot must complete/cancel prior methods)
   b. Drag and drop or browse for the .CSV file with liquid definitions and transfer steps.
   c. Check or modify the remaining runtime parameters. Notably the starting tip location (if the pipette box is not brand new) and the Left/Right location of the required P20 pipette.

**Starting Tip Rack:** This defines the location of the first 20 µL tip rack to be used. If Slot 1 is selected and its tips are exhausted during run, the OT-2 will automatically continue from Slot 9, which must therefore contain a full rack. If Slot 9 is chosen as the starting rack, non-full racks may be used; however, once Slot 9 is depleted, the system will not revert to Slot 1.

**Starting Tip Letter** and **Start Tip Number**: Location of the first row and column for the pipette tip box; The pipette picks up new tips by travel down each column, then shifting to the right. This feature is helpful if your tip rack is not full. If full, start with position A1.

**Pipette Location**: Choosing the Left/Right location of the required P20 pipette on the robot, from the perspective of someone standing in front of it.

**Other Pipette Choice** and **Other Pipette Location**: Select "None" or specify which additional pipette is already mounted and its location. The robot will not use the other pipette during this method.
**PCR Cycles**: Number of amplification cycles on the Thermocycler module.
**Elongation Minutes** and **Annealing Temperature**: Minutes to elongate at 72°C per PCR cycle, and the primer annealing temperature (°C) bind the template DNA.
**Temperature Module Cooling**: True (pause protocol until Temperature module cools to 14°C) or False (Cooling turned off for the entire run, pipetting starts immediately)

8. After confirming the runtime parameters, verify the final robot setup:
   a. Green checkmarks indicate if the pipettes and tips are calibrated and ready to use. If this data is not available, the Opentrons app will provide prompts to set this up.
   b. Run Labware Position Check to ensure the pipettes interact properly with labware. See **Supplementary Method 1**, follow the prompts built in the Opentrons App or this detailed online walk-through: Creating labware offsets with Labware Position Check
   c. A graphical overview of the labware and liquids allows the user to load and check that everything is in the correct position. If a run has 24 or fewer input samples (example below), tube_rack2 through tube_rack4 remain empty or can be omitted.

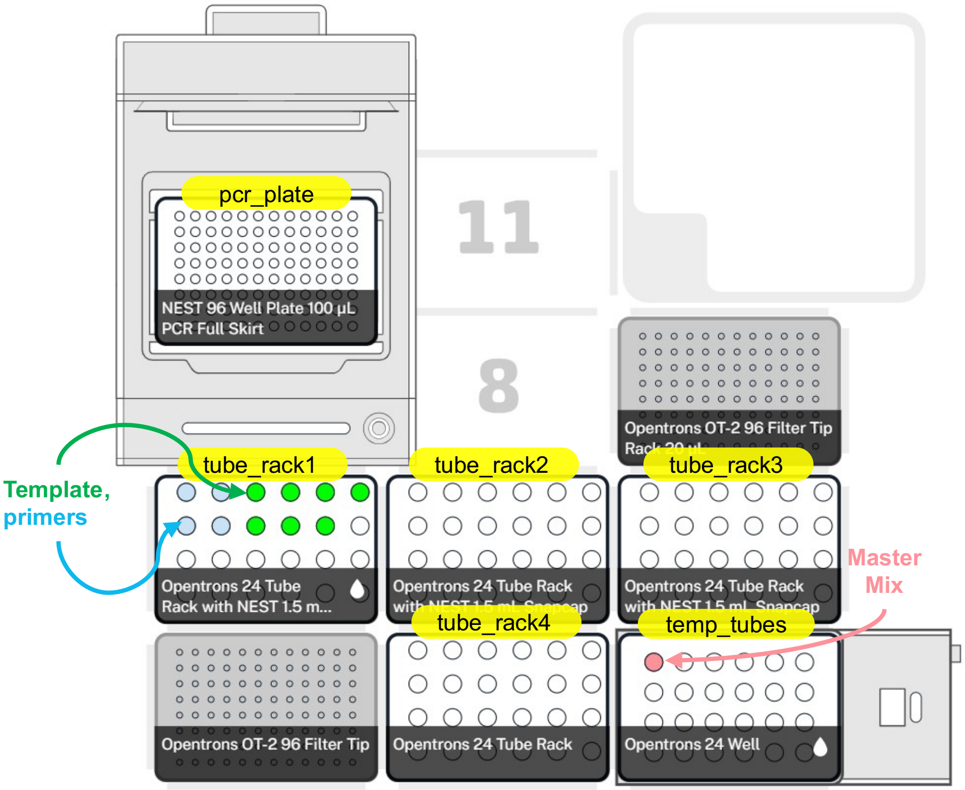

9. After checking the verification steps, click **Start Run** and enjoy the automated protocol.

### BOTany2B-PCR for Off-Deck Thermocycler

This protocol will set up PCR reactions in 8-well strips (the volume must be 200 µL; or alternatively, a 96-well PCR plate that can selected in the sample table) using components in 1.5 mL snap-cap tubes. The robot will add all the parts in the user-defined order. Compared to the **BOTany2A** protocol, the **BOTany2B** version does not require the on-deck Thermocycler and adds support for PCR strips that are compatible with many PCR machines found in molecular biology labs.

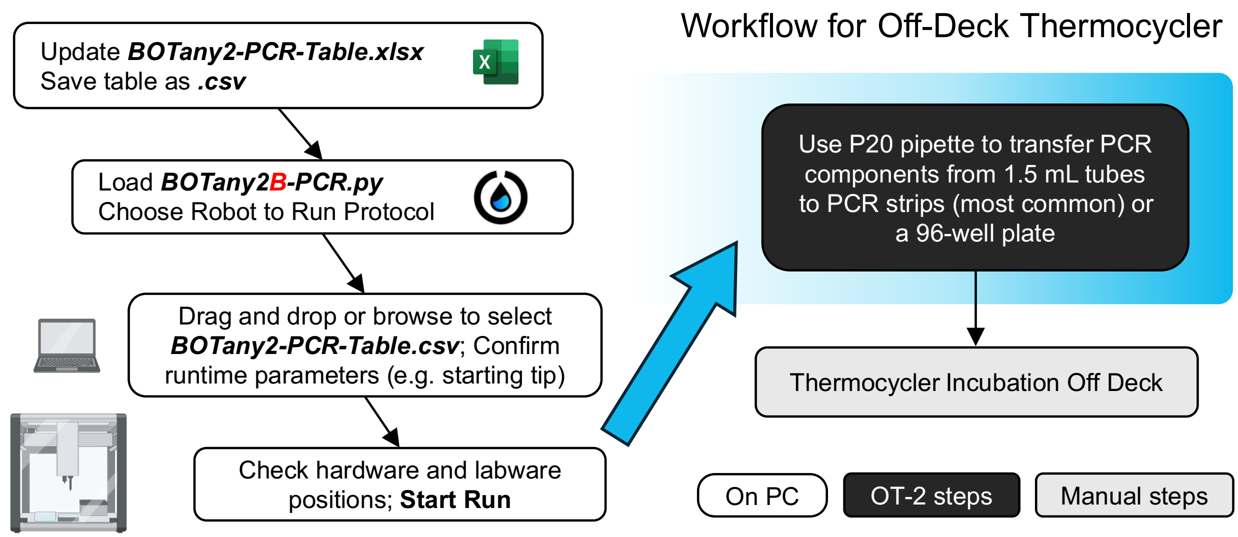

| **Required Hardware** | **API Name** |
|---|---|
| ○ P20 pipette | p20_single_gen2 |
| ○ Cooling block | temperature module gen2 |

| **Labware Quantity** | **API Name** |
|---|---|
| ○ 1, pre-chilled[2] | opentrons_24_aluminumblock_nest_1.5ml_snapcap |
| ○ 1 to 4 | opentrons_24_tuberack_nest_1.5ml_snapcap |
| ○ 1 to 12 | opentrons_96_aluminumblock_generic_pcr_strip_200µL |
| (Alternative: 1 plate) | opentrons_96_wellplate_200ul_pcr_full_skirt |

#### BOTany2B-PCR Instructions

1. Open the ***BOTany2-PCR-Table.xlsx*** template in Excel.
2. PCR parts are stored in 1.5 mL snap-cap tube racks (labelled tube_rack1 to tube_rack4). Each rack can hold up to 24 tubes (for up to 96 input parts). The temperature module (labelled temp_tubes) can chill up to 24 additional tubes at 14°C.
3. Edit the **Initial Liquid Definitions** (green headings) to specify which liquids will be loaded.
4. Edit **Pipetting Steps** (blue headings) to define volume transfers and robotic movements

[2] Pre-chilling aluminum blocks in a fridge or on ice can expedite the cooling of temperature module.

5. Save the updated Excel table in the .CSV format file [CSV UTF-8 format].
6. Open the Opentrons app. Import the ***BOTany2B-PCR.py*** script in the Protocols tab.

7. Click the three vertical dots on the top right of the protocol to **Start setup**.
   a. Choose Robot to set up runtime parameter (select from available wireless or wired connections; to become available, OT-2 robot must complete/cancel prior methods)
   b. Drag and drop or browse for the .CSV file with liquid definitions and transfer steps.
   c. Check or modify the remaining runtime parameters. Notably the starting tip location (if the pipette box is not brand new) and the Left/Right location of the required P20 pipette.

   **Temperature Module Cooling**: True (pause protocol until Temperature module cools to 14°C) or False (Cooling turned off for the entire run, pipetting starts immediately)

8. After confirming the runtime parameters, verify the final robot setup:
   a. Green checkmarks indicate if the pipettes and tips are calibrated and ready to use. If this data is not available, the Opentrons app will provide prompts to set this up.
   b. Run Labware Position Check to ensure the pipettes interact properly with labware. See **Supplementary Method 1**, follow the prompts built in the Opentrons App or this detailed online walk-through: Creating labware offsets with Labware Position Check
   c. A graphical overview of the labware and liquids allows the user to load and check that everything is in the correct position. If a run has 24 or fewer input samples (example below), tube_rack2 through tube_rack4 remain empty or can be omitted.

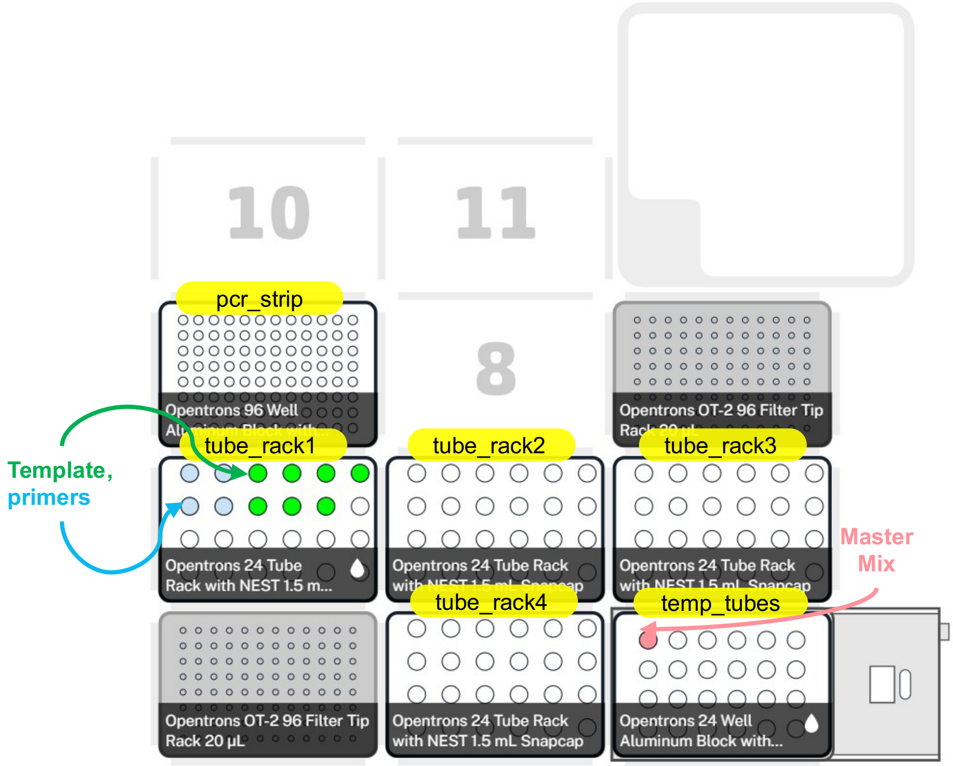

9. After checking the verification steps, click **Start Run** and manually move the PCR samples to an off-deck thermocycler for incubations when the OT-2 Run is complete.
