## Supplementary material for "BOTany Methods: Accessible Automation for Plant Synthetic Biology": Supplementary Method S3.pdf

---

### BOTany3A-MoClo for OT-2 Thermocycler

This protocol fully automates DNA assembly using Plant Modular Cloning (MoClo; or similar Golden Gate standards) in a 96-well plate using DNA components and enzyme mixtures loaded in 1.5 mL tubes. After pipetting the user-defined parts in the desired order, the robot will immediately run digestion and ligation cycles using the on-deck Opentrons Thermocycler.

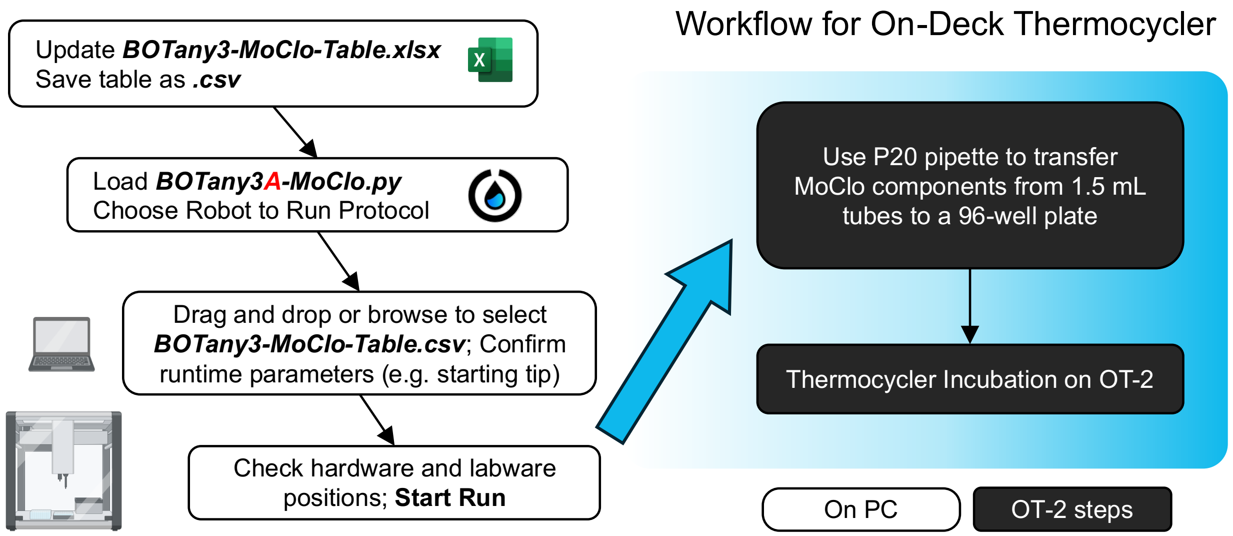

| **Required Hardware** | **API Name** |
|---|---|
| ○ P20 pipette | p20_single_gen2 |
| ○ Cooling block | temperature module gen2 |
| ○ Thermocycler | thermocyclerModuleV2 |

| **Labware Quantity** | **API Name** |
|---|---|
| ○ 1, pre-chilled[1] | opentrons_24_aluminumblock_nest_1.5ml_snapcap |
| ○ 1 to 4 | opentrons_24_tuberack_nest_1.5ml_snapcap |
| ○ 1 | nest_96_wellplate_100ul_pcr_full_skirt |

**Digest-Ligate Cycles**: Number of digestion-ligation cycles on the Thermocycler module.
**Digestion Minutes:** Minutes to hold at 37°C per digestion-ligation cycle.
**Ligation Minutes** and **Ligation Temperature**: Minutes and Temperature (°C) to ligate DNA during each cycle.

**Temperature Module Cooling**: True (pause protocol until Temperature module cools to 14°C) or False (Cooling turned off for the entire run, pipetting starts immediately)

8. After confirming the runtime parameters, verify the final robot setup:
   a. Green checkmarks indicate if the pipettes and tips are calibrated and ready to use. If this data is not available, the Opentrons app will provide prompts to set this up.
   b. Run Labware Position Check to ensure the pipettes interact properly with labware. See **Supplementary Method 1**, follow the prompts built in the Opentrons App or this detailed online walk-through: [Creating labware offsets with Labware Position Check]
   c. A graphical overview of the labware and liquids allows the user to load and check that everything is in the correct position. If a run has 24 or fewer input samples (example below), tube_rack2 through tube_rack4 remain empty or can be omitted.

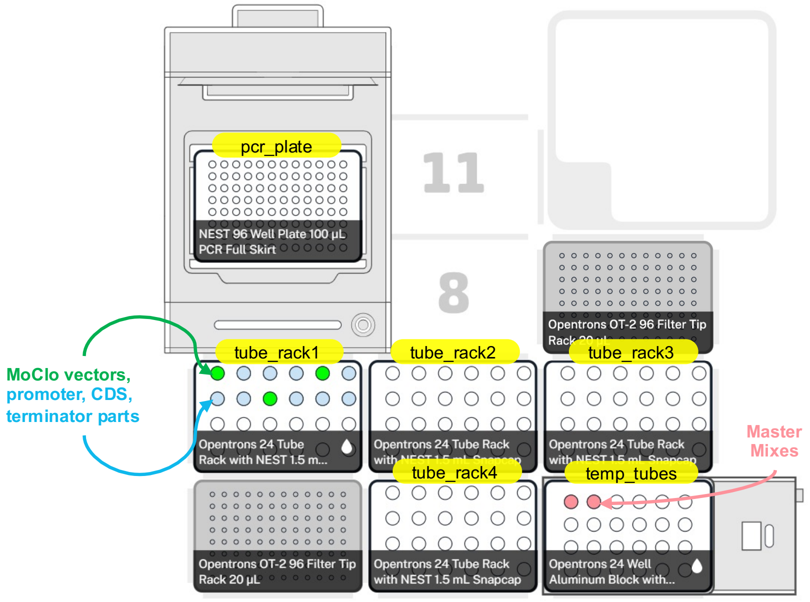

9. After checking the verification steps, click **Start Run** and enjoy the automated protocol.

### BOTany3B-MoClo for Off-Deck Thermocycler

This protocol allows DNA assembly reactions in 8-well PCR strips (the volume must be 200 µL; or alternatively, a 96-well PCR plate that can selected in the sample table) using components in 1.5 mL snap-cap tubes. The robot will add all the parts in user-defined order and volume from source wells to destination wells. Compared to the **BOTany3A** protocol, the **BOTany3B** version does not require the on-deck Thermocycler and adds support for PCR strips that are compatible with many PCR machines found in molecular biology labs.

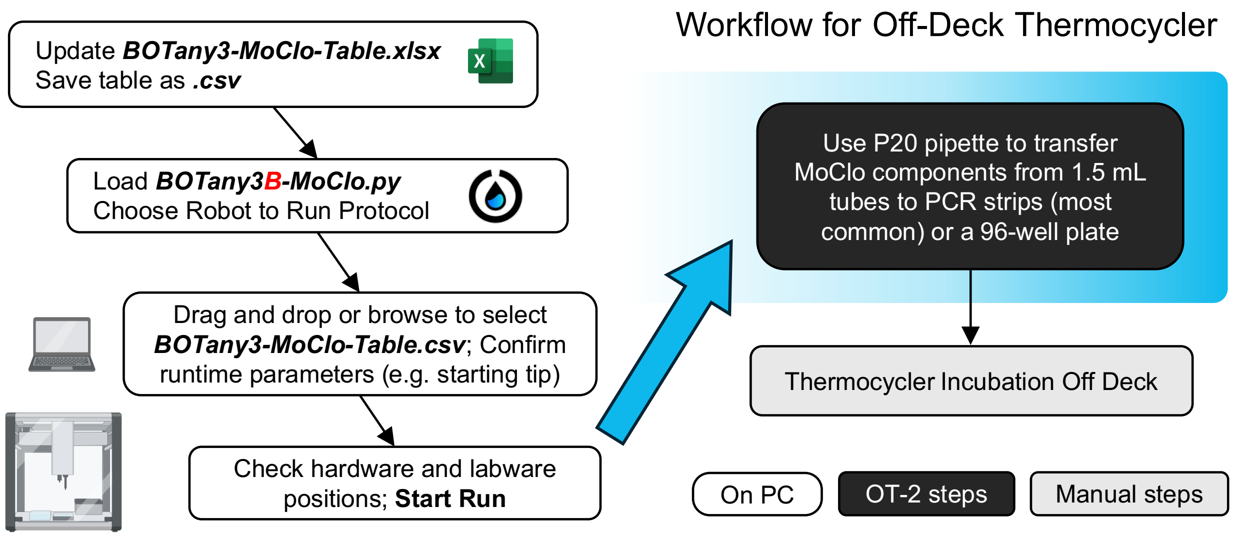

| **Required Hardware** | **API Name** |
|---|---|
| ○ P20 pipette | p20_single_gen2 |
| ○ Cooling block | temperature module gen2 |

| **Labware Quantity** | **API Name** |
|---|---|
| ○ 1, pre-chilled[2] | opentrons_24_aluminumblock_nest_1.5ml_snapcap |
| ○ 1 to 4 | opentrons_24_tuberack_nest_1.5ml_snapcap |
| ○ 1 to 12 | opentrons_96_aluminumblock_generic_pcr_strip_200µL |
| (Alternative: 1 plate) | nest_96_wellplate_100ul_pcr_full_skirt |

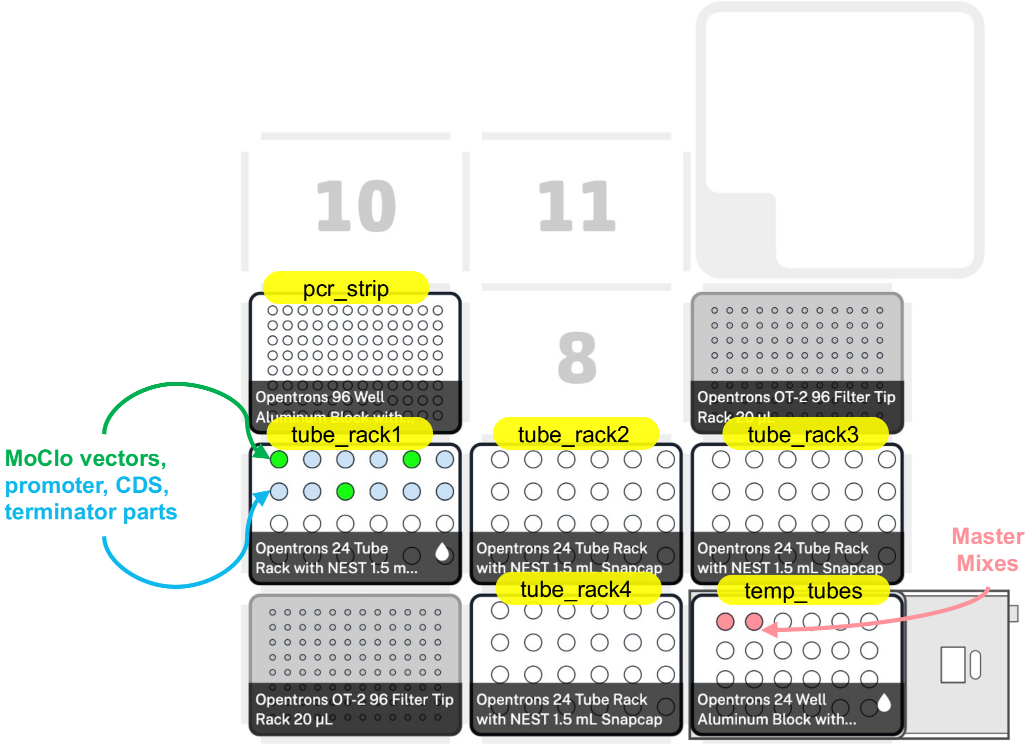

9. After checking the verification steps, click **Start Run** and manually move the PCR samples to an off-deck thermocycler for incubations when the OT-2 Run is complete.
