## Supplementary material for "BOTany Methods: Accessible Automation for Plant Synthetic Biology": Supplementary Method S4.pdf

### BOTany4-Shock&Go for Bacterial Transformation

This protocol automates *E. coli* cell transformation in a miniaturized PCR strip format (up to 96 samples). The robot will first add 20 µL of *E. coli* competent cells to sterile 200 µL PCR tubes, followed by 2 µL of DNA assembly mixture. Cells are heat shocked using the Opentrons Thermocycler Module GEN2 and automatically cooled. The OT-2 will add 150 µL outgrowth medium to aid cell recovery on the same module.

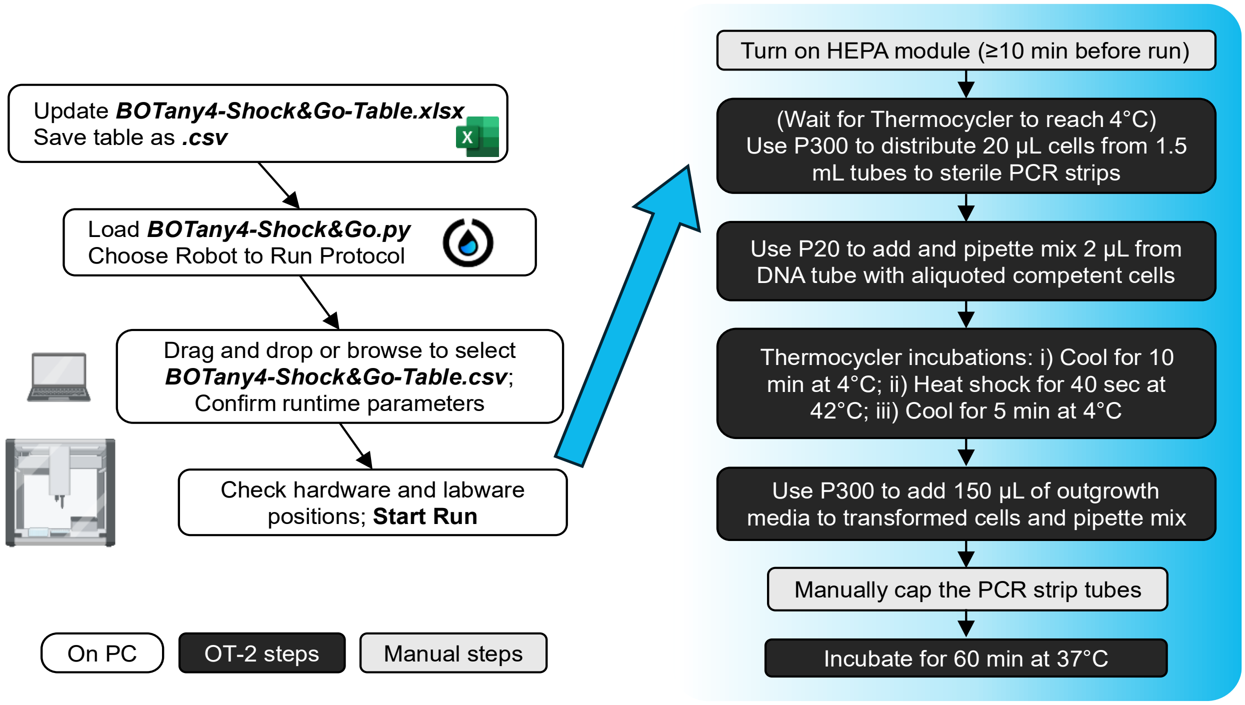


| **Required Hardware** | **API Name** |
|---|---|
| ○ P20 pipette | p20_single_gen2 |
| ○ P300 pipette | p300_single_gen2 |
| ○ Cooling block | temperature module gen2 |
| ○ Thermocycler | thermocyclerModuleV2 |
| ○ HEPA Module | *not applicable (manually controlled)* |

| **Labware Quantity** | **API Name** |
|---|---|
| ○ 1, pre-chilled[1] | opentrons_96_aluminumblock_generic_pcr_strip_200µL |
| ○ 1, pre-chilled | opentrons_24_aluminumblock_nest_1.5ml_snapcap |
| ○ 1 | opentrons_10_tuberack_falcon_4x50ml_6x15ml_conical |

#### BOTany4-Shock&Go Instructions

1. Open the ***BOTany4-Shock&Go.xlsx*** template in Excel.

[1] Pre-chill aluminum blocks in a fridge or on ice.

2. Edit the **Initial Liquid Definitions** (green headings) to specify which liquids will be loaded in the robot. *Note:* OT-2 cannot sense if liquids are loaded correctly or their volumes.

| | |
|---|---|
| **Labware** | **media_rack** or **cell_rack** or **DNA_tube** (case-sensitive names) |
| **Initial_Wells** | location of the liquid well/tube within a particular labware |
| **Initial_Volume** | the initial volume of said liquid in µL |
| **Liquid_Name** | any name you wish to assign the liquid |
| **Description** | any description you wish to assign the liquid (optional) |
| **Color** | HEX color codes that will show up in the OT-2 software deck view. |

3. Edit **Pipetting Steps** (**Cell Transfer**; blue headings) to define volume transfers:

| | |
|---|---|
| **Source_Labware** | case-sensitive name of the labware to aspirate (take) liquid from |
| **Source_Well** | position of well/tube within the Source_Labware |
| **Destination_Labware** | exact name of labware to dispense liquid into |
| **Destination_Well** | destination well/tube within the Destination_Labware |
| **Transfer_Volume** | liquid transfer volumes in microliters (µL) |
| **Pick_Up_Tip** | TRUE = new tips, FALSE = re-use tip (contamination risk later) |
| **Pipette_Choice** | Left/Right location of required pipette (when facing the OT-2 front) |

4. Edit **Pipetting Steps** (**DNA Transfer**; pink headings): same procedure as above but use new tips each sample to avoid cross-contamination and the DNA volume should not exceed 10% of the initial cell volume (e.g. 2 µL of Golden Gate reaction per 20 µL competent cell aliquot).
5. Edit **Pipetting Steps** (**Media Transfer**; orange headings): same procedure as above but use the initial tip can be reused when adding 150 µL of recovery media with the P300 pipette from a safe height.
6. Save the updated Excel table in the .CSV format file [CSV UTF-8 format].

7. Open the Opentrons app. Import the ***BOTany4-Shock&Go.py*** script in the Protocols tab.
8. Manually turn on the HEPA module (max flow) for >10 min with the OT-2 door closed.

9. Click the three vertical dots on the top right of the protocol to **Start setup**.
   a. Choose Robot to Run protocol (select from available wireless or wired connections; to become available, OT-2 robot must complete/cancel prior methods)
   b. Drag and drop or browse for the .CSV file with liquid definitions and transfer steps.
   c. Check or modify the final runtime parameters. Notably the starting tip location (if the pipette box is not brand new) and the Left/Right location of the pipettes.

**Starting Tip Letter** and **Start Tip Number**: Location of the first row and column for the pipette tip box for either the left or right pipette; Each pipette picks up new tips by traveling down the column, then shifting to the right. This feature is helpful if your tip rack is not full. If full, start with position A1.

**Pipette Location**: Choosing the Left/Right location of the required P20 and P300 pipettes on the robot, from the perspective of someone standing in front of it.

10. After confirming the runtime parameters, verify the final robot setup:
    a. Green checkmarks indicate if the pipettes and tips are calibrated and ready to use. If this data is not available, the Opentrons app will provide prompts to set this up.
    b. Run Labware Position Check to ensure the pipettes interact properly with labware. See **Supplementary Method 1**, follow the prompts built in the Opentrons App or this detailed online walk-through: Creating labware offsets with Labware Position Check
    c. A graphical overview of the labware and liquids allows the user to load and check that everything is in the correct position.

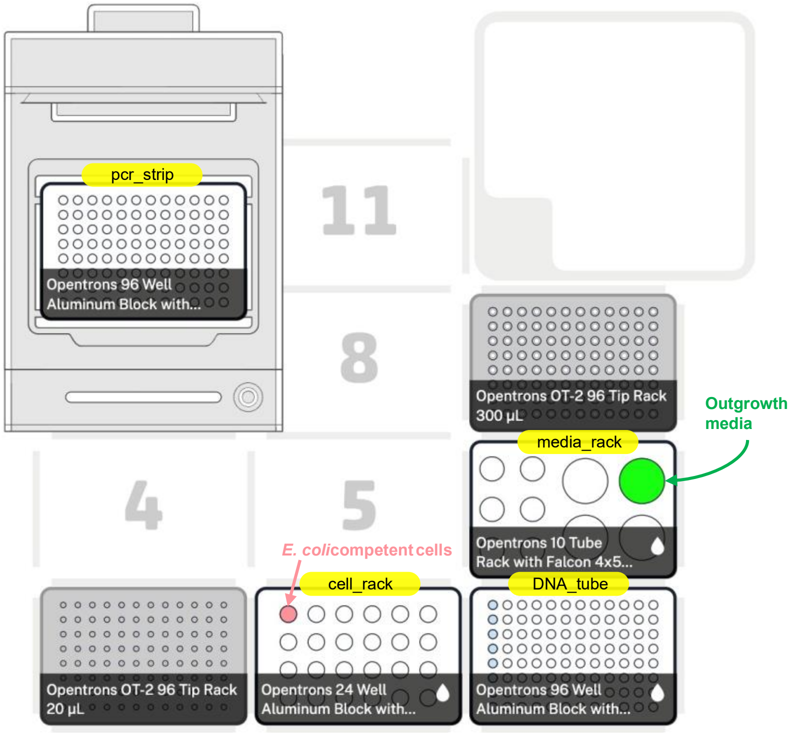


11. After checking the verification steps, click **Start Run** and all steps will run automatically until each tube has outgrowth media.
12. A prompt will appear at this point to instruct users to cap the lids to avoid evaporation during the next 1 h outgrowth. After capping lids, HEPA module can be turned off.

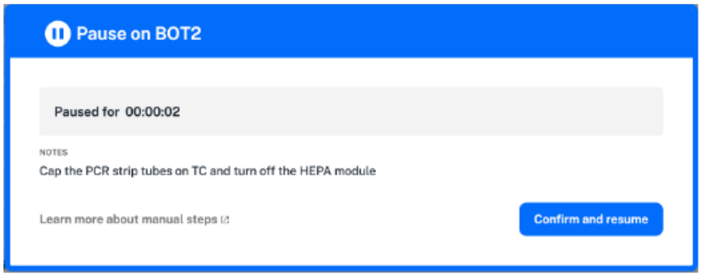


13. After the outgrowth, pipette mix and plate at least 50 µL cell cultures on LB agar plates with proper antibiotics.
