## Supplementary material for "BOTany Methods: Accessible Automation for Plant Synthetic Biology": Supplementary Method S5.pdf

### BOTany5-MagBead Plasmid Extraction

This protocol allows plasmid extraction using Zymo Research's MagBead plasmid extraction kit from 8-96 samples. The first step is to manually get clear lysate and transfer to a Zymo collection plate. The second step is to bind, wash, and elute plasmid DNA on OT-2 using 8-channel 300 pipette.

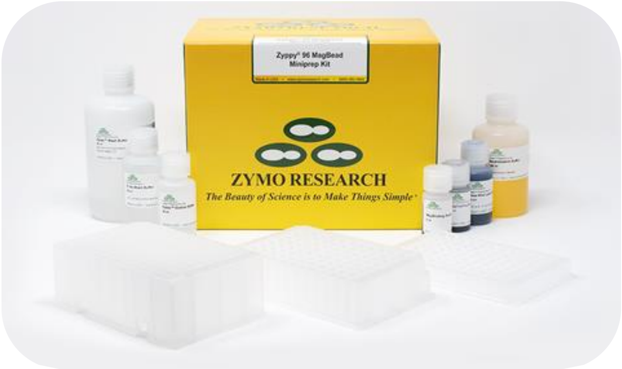


**Required Kit**: [Zyppy-96 Plasmid MagBead Kit (Cat # D4100 Zymo Research)]

#### BOTany5-MagBead Manual Instructions

| **Required Hardware** | **Brand & Cat #** |
|---|---|
| Magnetic stand | Zymo Research P1005 |

| **Labware Quantity** | **Labware name** | **Brand & Cat #** |
|---|---|---|
| o 1 to 4 | 24 deep well plate | Sigma-Aldrich AXYPDW10ML24C |
| o 1 to 4 | Plate sealer | Sigma-Aldrich Z763624-100EA |
| o 1 | Zymo Collection Plate | Zymo Research C2002 |

Bacterial lysis is manually performed in 24 deep well plate format, off-deck:

1. Grow single colonies of transgenic *E. coli* in 1.6 mL LB medium supplemented with proper antibiotic in 24 deep well plate overnight at 37 °C. Use 800 µL *E. coli* culture to make glycerol stock.
2. Add 100 µL Blue Lysis Buffer directly to each well with remaining (approximately 750 µL) *E. coli* culture. Gently swirl the plate and incubate at room temperature for 1 min. The sample should turn blue.
3. Add 450 µL Neutralization Buffer to each well. Gently swirl the solution until it turns yellow.
4. Vortex mix the Clearing Beads Buffer bottle to resuspend the beads. Add 50 µL Clearing Beads Buffer to each well. The sample will turn brown color when the binding is complete.
5. Place the 24 deep well plate on the magnetic stand (sold separately from the kit) and wait 3-5 min until beads separate to give a clear lysate.
6. Transfer 650 µL cleared lysates to a Zymo Collection Plate (supplied).

#### BOTany5-MagBead OT-2 Instructions

| **Required Hardware** | **API Name** |
|---|---|
| ○ 8-channel P300 | p300_multi_gen2 |
| ○ Heater Shaker Module | heaterShakerModuleV1 |
| ○ Magnetic Module Gen2 | mag_mod |

| **Labware Quantity** | **API Name** |
|---|---|
| ○ 1 | nest_12_reservoir_15ml |
| ○ 1 | nest_1_reservoir_195ml |
| ○ 1 | nest_96_wellplate_2ml_deep |
| ○ 1 | zymocollection_96_wellplate_1200ul |
| ○ 1 | zymoelution_96_wellplate_90ul |

**Note**: The last two items are plates from the Zymo kit and require custom .json files.

1. Open the Opentrons app. Import the ***BOTany5-Magbead.py*** script in the Protocols tab.
2. Click the three vertical dots on the top right of the protocol to **Start setup**.
   a. Choose Robot to set up runtime parameters (select from available wireless or wired connections; to become available, OT-2 robot must complete/cancel prior methods)
   b. Check and modify the final runtime parameters:
      **Number of samples**: number of clear lysates loaded on the collection plate. This number should be the multiples of 8.
      **Sample Starting Column**: pick the starting column where clear lysates were loaded on the collection plate (e.g. 1 = $1^{st}$ column).
      **Settling Time:** time in min for magnetic beads to settle on the module.
      **Elution Plate:** type of plate used as elution plate. Options include:
         opentrons_96_wellplate_200µL_pcr_full_skirt
         nest_96_wellplate_100µL_pcr_full_skirt
         zymoelution_96_wellplate_90µL (requires custom .json file)
      **Collection Plate:** Use the zymocollection_96_wellplate_1200µL (requires custom .json file).
      **Fast run:** choosing this option will skip the second Zyppy buffer wash and shorten incubations on magnetic module to 6 sec. This feature is helpful for testing the protocol with water.
      **8-Channel P300 Location:** Left/Right mount location of the required pipette.

3. Click the three vertical dots on the top right of the protocol to **Start setup**.
4. After confirming the runtime parameters, verify the final robot setup:
   a. Green checkmarks indicate if the pipettes and tips are calibrated and ready to use. If this data is not available, the Opentrons app will provide prompts to set this up.
   b. Run Labware Position Check to ensure the pipettes interact properly with labware. See **Supplementary Method 1**, follow the prompts built in the Opentrons App or this detailed online walk-through: [Creating labware offsets with Labware Position Check]
   c. A graphical overview of the labware and liquids allows the user to load and check that everything is in the correct position.

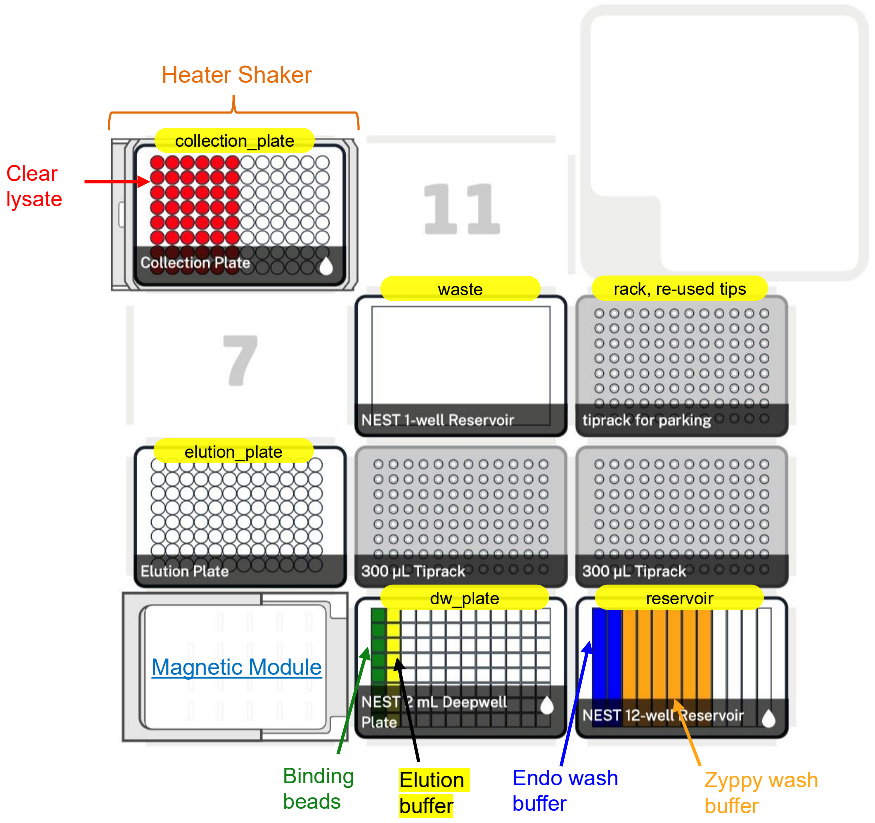


#### OT-2 On-Deck Procedures

OT-2 will carry out the following steps in a 96-well format with [**Manual**] interventions. Unless otherwise noted, all interventions refer to **Zymo Collection Plate**.

1. Mix the binding buffer 5 times in the first column of NEST 96 Deep Well Plate (slot 7). Add 30 µL binding bead buffer to the Zymo Collection Plate on Heater Shaker.
2. Vortex Zymo Collection Plate on Heater Shaker at 1000 rpm for 5 min.

3. **[Manual]** Move the Zymo Collection Plate: Heater Shaker → Magnetic Module.
4. Engage magnets to the height of 3.9 mm. Let beads settle for 1 min.
5. Aspirate 650 µL liquid from each well of Zymo Collection Plate and discard it into the waste reservoir (slot 8). Park the tips back on the rack (slot 9) after discarding the waste.

6. **[Manual]** Disengage magnets and move plate: Magnetic Module → Heater Shaker.
7. Distribute 200 µL Endo wash buffer (slot 11, column 1-2) to each well.
8. Vortex Zymo Collection Plate on the Heater Shaker at 1100 rpm for 90 sec.

9. **[Manual]** Move the Zymo Collection Plate: Heater Shaker → Magnetic Module
   Engage magnets to the height of 3.9 mm. Let beads settle for 1 min.
   Aspirate 200 µL from each well of the Zymo plate and discard it into the waste reservoir (slot 8). Park the tips back on the rack (slot 9) after discarding the waste.

10. **[Manual]** Disengage magnets and move the plate: Magnetic Module → Heater Shaker.
11. Distribute 300 µL Zyppy wash buffer (slot 11, column 3-7) to each well.

12. Vortex the Zymo Collection Plate on Heater Shaker at 1200 rpm for 90 sec.

13. **[Manual]** Move the Zymo Collection Plate: Heater Shaker → Magnetic Module
    Engage magnets to the height of 3.9 mm. Let beads settle for 1 min.
    Aspirate 300 µL from each well of Zymo Collection Plate and discard it into the waste reservoir (slot 8). Park the tips back on the rack (slot 9) after discarding the waste.

14. **[Manual]** Disengage magnets and move plate: Magnetic Module → Heater Shaker.
15. Perform the Zyppy wash step a second time by repeating step 11 through 14.

16. **[Manual]** Turn on the HEPA module to accelerate the sample drying process.
17. Vortex Zymo Collection Plate on the Heater Shaker at 1800 rpm at 65 C for 10 min.
18. Turn off the HEPA module manually and deactivate the Heater Shaker.
19. Distribute 40 µL Elution buffer (slot 7, column 2) to each well of Zymo Collection Plate.
20. Vortex Zymo Collection Plate on the Heater Shaker at 1200 rpm for 90 sec.

21. **[Manual]** Move the Zymo Collection Plate: Heater Shaker → Magnetic Module
    Engage magnets to the height of 3.9 mm. Let beads settle for 1 min.

22. Transfer 30 µL liquid from each well of Zymo Collection Plate (from 3 mm above the bottom) into the Zymo Elution Plate (slot 4).

*Optional step*: If beads are visible in the elution plate before DNA quantification, they can be pelleted with the Zymo MagStand. Beads do not generally interfere with plasmid use.
