## Supplementary material for "BOTany Methods: Accessible Automation for Plant Synthetic Biology": Supplementary Method S6.pdf

### BOTany6-Universal Protocol

This protocol enables universal volume transfer on OT-2 using single channel pipettes.

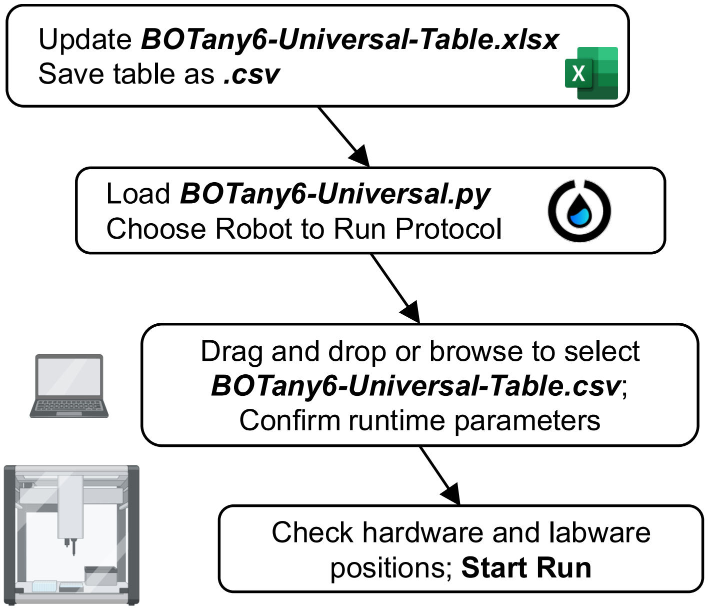


**Hardware**
- User-defined from several pre-configured options

**Labware**
- User-defined from several pre-configured options

#### BOTany6-Universal Instructions

1. Open the ***BOTany6-Universal-Table.xlsx***, and update the colored sections:
   a. **Tip Rack Definition**
   **Rack_Name:** unique name you want to give the tip rack. This is decided by the user.
   **Rack_API:** case-sensitive labware API name (defined by Opentrons)
   **Rack_Location:** location of the slot the tip rack on the OT-2 deck, from 1-11.

   b. **On-Deck Labware Definition** (for labware placed directly on OT-2 deck)
   **Labware_Name**: whatever unique name you want to give the labware.
   **Labware_API:** API id from Opentrons.
   **Labware_Location:** location of the slot the labware is in on the OT-2, from 1-11.

   c. Module Definition (e.g. Opentrons Thermocycler, Heater-Shaker, etc.)
   **Module_Name**: whatever unique name you want to give the module.
   **Module_API:** API id from Opentrons.

**Module_Location**: location of the slot the module is in on the OT-2, from 1-11. If to use Thermocycler module, the location should be “None”.

d. On-Module Labware Definition (mounted on top of modules)
**Base_Labware_Name**: exact name of a module from step **1c**
**Top_Labware_Name**: any desired name for the labware loaded on top of the module.
**Top_Labware_API**: case-sensitive API name from Opentrons.

e. **Initial Liquid Definitions:** to specify which liquids will be loaded in the robot. *Note:* OT-2 cannot sense if liquids are loaded correctly or their volumes.
**Labware**: exact name of labware defined in earlier steps
**Initial_Wells**: location of the liquid well/tube within a particular labware
**Initial_Volume**: the initial volume of said liquid in μL
**Liquid_Name**: any name you wish to assign the liquid
**Description**: description you wish to assign the liquid (optional)
**Color:** HEX color codes that will show up in the OT-2 software deck view.

f. **Pipetting Steps**: to define volume transfers:

| | |
|---|---|
| **Source_Labware** | case-sensitive name of the labware to aspirate (take) liquid from |
| **Source_Well** | position of well/tube within the Source_Labware |
| **Destination_Labware** | exact name of labware to dispense liquid into |
| **Destination_Well** | destination well/tube within the Destination_Labware |
| **Transfer_Volume** | liquid transfer volumes in microliters (μL) |
| **Pick_Up_Tip** | <u>TRUE</u> = new tips, <u>FALSE</u> = re-use tip (contamination risk later) |
| **Pipette_Choice** | Left/Right location of required pipette (when facing the OT-2 front) |

g. Save the updated Excel table in the .CSV format file [CSV UTF-8 format].

2. Open the Opentrons app. Import the ***BOTany6-Universal.py*** script in the Protocols tab.

3. Click the three vertical dots on the top right of the protocol to Start setup.
   a. Choose Robot to Run protocol (select from available wireless or wired connections; to become available, OT-2 robot must complete/cancel prior methods)
   b. Drag and drop or browse for the .CSV file with liquid definitions and transfer steps.
   c. Check or modify the final runtime parameters, such as the starting location of the tips and selected pipettes.

**Left/Right Pip Start Tip Letter** and **Number:** Location of the first row and column for the pipette tip box for either the left or right pipette; Each pipette picks up new tips by traveling down the column, then shifting to the right. This feature is helpful if your tip rack is not full. If full, start with position A1.

**Left/Right Pipette Choice**: Choosing the Left/Right location of the desired pipettes, from the perspective of someone standing in front of the OT-2. Note that 8-channel pipettes can be mounted but cannot be used for this protocol.

4. After confirming the runtime parameters, verify the final robot setup:
   a. Green checkmarks indicate if the pipettes and tips are calibrated and ready to use. If this data is not available, the Opentrons app will provide prompts to set this up.
   b. Run Labware Position Check to ensure the pipettes interact properly with labware. See **Supplementary Method 1**, follow the prompts built in the Opentrons App or this detailed online walk-through: Creating labware offsets with Labware Position Check
   c. A graphical overview of the labware and liquids allows the user to load and check that everything is in the correct position.

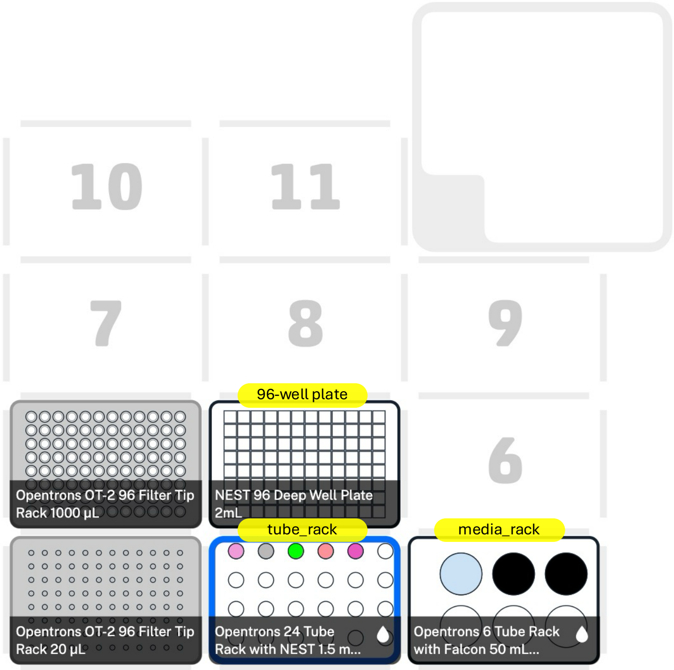


This deck layout represents the example in Fig. 4 and is also reflected in *BOTany6-Universal-Table.xlsx*.

5. After checking the verification steps, click **Start Run** and enjoy the automated protocol. Tip: a new protocol can be tested with water or even without liquids to check if the OT-2 robot follows the desired instructions.

#### Optional Steps for Further Customization

1. Custom Labware:
   a. Custom labware can be created via the Opentrons Labware Creator. This labware can then be imported into the OT-2 app, and it's API can be used like any other labware: https://labware.opentrons.com/#/create
2. Liquid Transfer Parameters:
   a. Aspiration and Dispensing - to modify the heights, speed, etc. of dispensing and aspirating liquid, follow the OT-2 manual for the lines curr_pip.aspirate and curr_pip.dispense: https://docs.opentrons.com/v2/basic_commands/liquids.html#liquid-control
