## Supplementary material for "BOTany Methods: Accessible Automation for Plant Synthetic Biology": Supplementary Tables and Figures.pdf

**Supplementary Table S1**. Summary of BOTany Excel and Python files on GitHub.

| Method name | Excel table name | Python file name |
|---|---|---|
| BOTany1-Primers | BOTany1-Primers.xlsx / .csv | BOTany1-Primers.py |
| BOTany2-PCR | BOTany2-PCR.xlsx / .csv | BOTany2A-PCR.py<br>BOTany2B-PCR.py |
| BOTany3-MoClo | BOTany3-MoClo.xlsx / .csv | BOTany3A-MoClo.py<br>BOTany3B-MoClo.py |
| BOTany4-Shock&Go | BOTany4-Shock&Go.xlsx / .csv | BOTany4-Shock&Go.py |
| BOTany5-MagBead | BOTany5-MagBead.xlsx / .csv | BOTany5-MagBead.py |
| BOTany6-Universal | BOTany6-Universal.xlsx / .csv | BOTany6-Universal.py |

**Supplementary Table S2**. Plasticware used in BOTany methods.

| Protocol used | Generic name | Opentrons API name | Brand | Cat# |
|---|---|---|---|---|
| BOTany1-Primers | 50 mL Falcon tube | opentrons_10_tuberack_falcon_4x50ml_6x15ml_conical | Sarstedt | 62.547.205 |
| | 2 mL screw cap tube | opentrons_24_tuberack_generic_2ml_screwcap | Sarstedt | 72.664 |
| | 1.5 mL snap cap tube | opentrons_24_tuberack_nest_1.5ml_snapcap | Sarstedt | 72.690.550 |
| BOTany2-PCR or BOTany3-MoClo | 200 µL PCR strip (BOTany 2B and 3B) | opentrons_96_aluminumblock_generic_pcr_strip_200ul | Sarstedt | 72.985.002 |
| | 100 µL PCR plate (BOTany2A and 3A) | nest_96_wellplate_100ul_pcr_full_skirt | Nest | 999-00050 |
| | 1.5 mL snap cap tube | opentrons_24_tuberack_nest_1.5ml_snapcap | Sarstedt | 72.690.550 |
| | | opentrons_24_aluminumblock_nest_1.5ml_snapcap | Sarstedt | 72.690.550 |
| BOTany4-Shock&Go | 200 µL PCR strip | opentrons_96_aluminumblock_generic_pcr_strip_200ul | Sarstedt | 72.985.002 |
| | 1.5 mL snap cap tube | opentrons_24_aluminumblock_nest_1.5ml_snapcap | Sarstedt | 72.690.550 |
| | 50 mL Falcon tube | opentrons_10_tuberack_falcon_4x50ml_6x15ml_conical | Sarstedt | 62.547.205 |
| BOTany5-MagBead | 12-well reservoir | nest_12_reservoir_15ml | Nest | 999-00076 |
| | 1-well reservoir | nest_1_reservoir_195ml | Nest | 999-00078 |
| | 2 mL deep well plate | NEST 2 mL Deepwell Plate | Nest | 999-00103 |
| | Zymo collection plate | zymocollection_96_wellplate_1200ul | Zymo Research | C2002 |
| | Zymo elution plate | zymoelution_96_wellplate_90ul | Zymo Research | C2003 |
| BOTany6-Universal | Customizable | Any Opentrons parts and custom labware | any | - |

**Supplementary Table S3**. Yield and purity of MagBead-extracted plasmids. Parameters were exported for NanoDrop.

| Plasmid | DNA (ng/µL) | A260/A280 | A260/A230 | A260 | A280 |
|---|---|---|---|---|---|
| E1 | 60.8 | 1.99 | 1.13 | 1.22 | 0.61 |
| F1 | 32.2 | 2.19 | 0.75 | 0.64 | 0.29 |
| G1 | 48.8 | 2.05 | 0.97 | 0.98 | 0.48 |
| H1 | 39.8 | 2.13 | 0.69 | 0.80 | 0.37 |
| A3 | 29.2 | 1.76 | 0.96 | 0.58 | 0.33 |
| B3 | 27.5 | 1.93 | 1.13 | 0.55 | 0.28 |
| C3 | 28.4 | 1.80 | 0.98 | 0.57 | 0.32 |
| D3 | 74.8 | 0.83 | 0.84 | 1.50 | 1.80 |
| E3 | 21.5 | 1.75 | 0.53 | 0.43 | 0.24 |
| F3 | 28.2 | 1.67 | 0.91 | 0.56 | 0.34 |
| G3 | 28.5 | 1.83 | 0.70 | 0.57 | 0.31 |
| H3 | 27.9 | 1.82 | 0.99 | 0.56 | 0.31 |

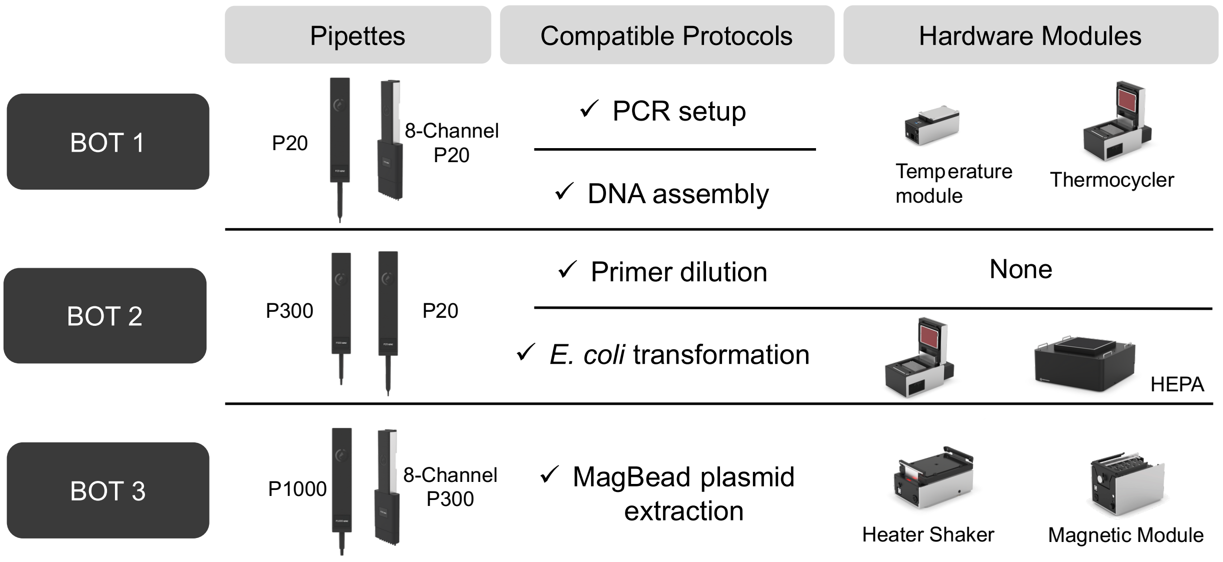


**Supplementary Figure S1.** BOTany Foundry Configuration. Example configuration with three separate robots that could be designated to carry out parallel methods described in this study. Alternatively, a single robot can perform any of the protocols if the hardware and labware is changed and calibrated between experimental runs.

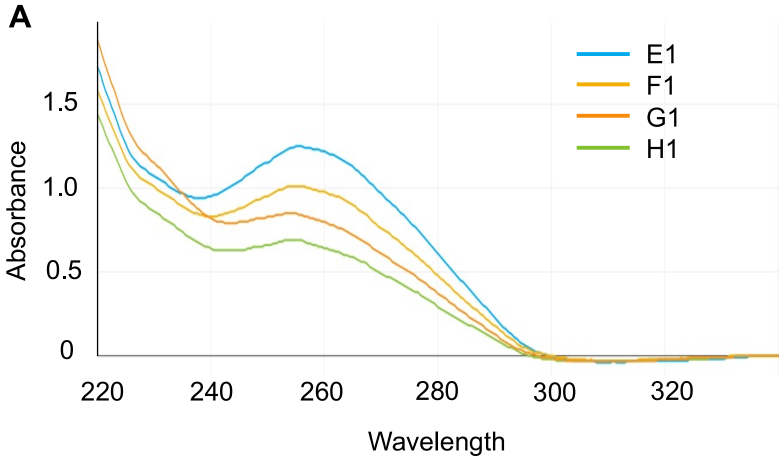


| ID | ng/μL | A260/A280 |
|---|---|---|
| E1 | 60.8 | 1.99 |
| F1 | 32.2 | 2.19 |
| G1 | 48.8 | 2.05 |
| H1 | 39.8 | 2.13 |

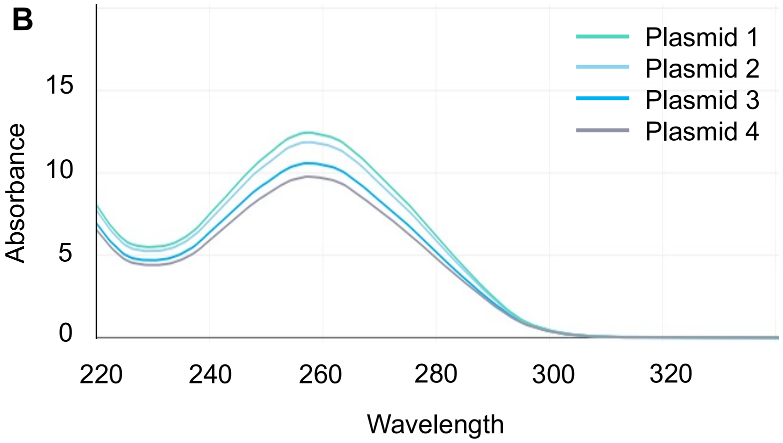


| ID | ng/μL | A260/A280 |
|---|---|---|
| 1 | 524.8 | 2.02 |
| 2 | 587.7 | 2.00 |
| 3 | 615.9 | 2.00 |
| 4 | 484.2 | 2.01 |

**Supplementary Figure S2**. Plasmids extracted with MagBead versus spin columns. NanoDrop spectra of plasmids from extracted using the MagBead method on the OT-2 **(A)** or manually using spin columns and centrifugation **(B)**. Quantity and quality of plasmids were monitored for 1 μL aliquots after DNA elution in 30 μL of HyClone water.

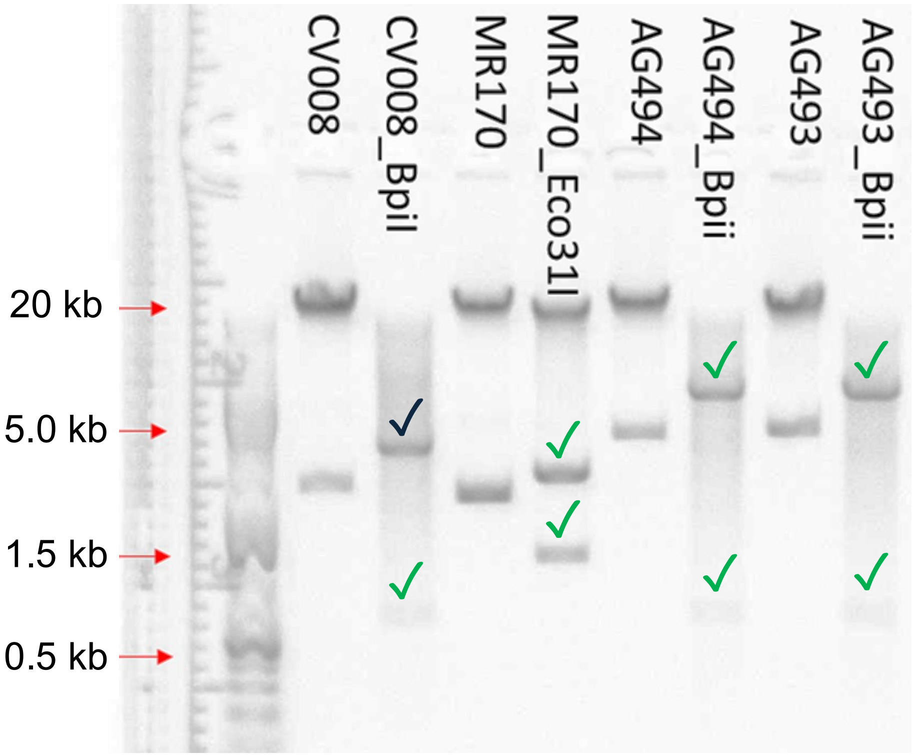


| Plasmid | Size in kb | Restriction Enzyme | Fragment 1 in kb | Fragment 2 in kb |
|---|---|---|---|---|
| CV008 | 3.3 | BpiI | 0.6 | 1.1 |
| MR170 | 3.0 | Eco31I | 1.0 | 2.1 |
| AG494 | 5.3 | BpiI | 0.6 | 4.7 |
| AG493 | 5.7 | BpiI | 0.6 | 5.1 |

**Supplementary Figure S3**. Test digestions of MagBead-extracted plasmids. The agarose gel image shows undigested and digested samples for four different plasmids, ranging from 3.0 to 5.7 kb (total size). Except for the CV008 plasmid which may have been linearized but not efficiently cut twice, plasmids showed the two expected sizes after digestion.  Large molecular weight DNA was also visible in the undigested controls.
